# Cross-attractor repertoire provides new perspective on structure-function relationship in the brain

**DOI:** 10.1101/2020.05.14.097196

**Authors:** Mengsen Zhang, Yinming Sun, Manish Saggar

## Abstract

The brain is a complex system exhibiting ever-evolving activity patterns without any external inputs or tasks. Such intrinsic dynamics (or lack thereof) are thought to play crucial roles in typical as well as atypical cognitive functioning. Linking the ever-changing intrinsic dynamics to the rather static anatomy is a challenging endeavor. Dynamical systems models are important tools for understanding how structure and function are linked in the brain. Here, we provide a novel modeling framework to examine such structure-function relations. Our deterministic approach complements previous modeling frameworks, which typically focus on noise-driven (or stochastic) dynamics near a single attractor. We examine the overall organizations of and coordination between all putative attractors. Using our approach, we first provide evidence that examining cross-attractor coordination between brain regions could better predict human functional connectivity than examining noise-driven near-attractor dynamics. Further, we observed that structural connections across scales modulate the energy costs of such cross-attractor coordination. Overall, our work provides a systematic framework for characterizing intrinsic brain dynamics as a web of cross-attractor transitions and associated energy costs. The framework may be used to predict transitions and energy costs associated with experimental or clinical interventions.

## 1 Introduction

A fundamental goal of neuroscience is to understand how the structure of the brain constrains its function [1]. The advent of neuroimaging techniques has enabled detailed, quantitative examination of the structure-function relation, often by comparing the structural and functional connectivity between brain regions [2, 3]. The resting-state functional connectivity is of particular interest, given its relevance to a variety of cognitive functions, neurological diseases, and psychiatric disorders [3, 4]. Though structural connectivity and functional connectivity are linearly correlated, the former does not entirely predict the latter—strong functional coupling exists between regions with only weak or indirect structural connections [2, 5]. Nonlinear dynamical models have been used to provide a mechanistic understanding of the structure-function relation [6–8] and provided many insights (e.g. [9–16]). While successful, previous dynamical system approaches open focus on dynamics near a single stable state, i.e., an attractor. However, biological systems such as the brain are often multistable [17–19], i.e., multiple attractors can coexist in the brain’s dynamical landscape. Such multistability begs the question of whether examining the overall layout of brain’s attractor states could better inform or complement what we know about the structure-function relation than explorations around a single state.

Intrinsic brain dynamics have long been observed [20, 21] but often treated as a baseline subtracted from task-positive activities. This baseline, however, is more active than meets the eye: it consumes the largest fraction of the brain’s energy resources, while task-related consumption adds little [22]. It constrains task performance and related neural activities across multiple time scales [23–25], and sustains alteration in neurological and psychiatric disorders [26–30]. In contrast to the restless dynamics is the (relatively) static structure—the anatomical connections between brain regions, which can be estimated non-invasively using large-scale tractography on diffusion-weighted images [31]. How can one compare the ever-changing with the unchanging? From a statistical perspective, one may compute the time-averaged features of the dynamics, such as the correlation between signals generated by two brain regions across time—a common measure of functional connectivity. Such functional connectivity patterns can be directly compared to structural ones through linear correlation [2]. From a dynamics perspective [6, 8, 32], the strength of anatomical connections can be incorporated as constant parameters in a system of differential equations, i.e. a dynamical system. The dynamical system, in turn, describes how the state of a model brain, endowed with realistic anatomy, would evolve over time. The time series generated by the model brain and the derived functional connectivity patterns can then be fitted to that of the real brain. Thus, a dynamical system naturally bridges between the unchanging structure and the ever-changing dynamics.

The brain’s nonlinearity gives rise to multistability and sustained oscillation [17–19, 33–36]. In a multistable landscape, multiple attractors coexist. That is, a brain may persistent in one of many qualitatively distinct patterns of activity depending on the environment and intrinsic factors. One popular modeling approach is to simulate the noise-driven dynamics near a chosen attractor, such as the low activity ground state, and compare it to the human resting brain dynamics (see [37] for a summary of different approaches). Such noise-driven exploration of a single attractor has been shown to exhibit key features of human resting brain dynamics, especially near criticality (e.g. [10, 12, 16]). On the other hand, noise-driven exploration beyond a single attractor—across multiple attractors or “ghost” attractors—has been shown to capture non-stationary resting brain dynamics and the switching between different dynamic functional connectivity patterns [11, 14, 15, 38]. The best fit to empirical data is often found near the onset of multistability [14, 38]. These observations suggest that examining the layout of the attractor repertoire over the entire multistable landscape could be crucial for understanding the organization of resting brain dynamics (c.f. [14]).

Complementing existing single-attractor approaches, the present work focuses on the deterministic features of the multistable landscape and examines their empirical relevance. Specifically, we systematically study the organization of the attractor repertoire as a window into the overall shape of the dynamic landscape. To do so, we use a biophysical network model that formally combines the reduced Wong-Wang model [12, 13, 39] and the Wilson-Cowan model [40, 41]. We first should that the model exhibits extensive multistability, i.e. a large repertoire of attractors to serve as landmarks of the landscape. Further, the model also allows us to examine, computationally and analytically, how structural features across scales shape this repertoire. It is important to note that here we used a broader definition structural features and not only include large-scale structural connectivity between brain regions, but also local recurrent connectivity within regions, and biophysical constraints at the cellular level. Using this modeling framework and a small dataset from the Human Connectome Project (HCP; n=100), we provide evidence with regards to how the cross-attractor relations in the repertoire could better capture key features of human resting functional connectivity and how such features are shaped by structural features across scales. Finally, we provide a novel framework to analyze the energy constraints for such cross-attractor coordination across different local and global structures.

## 2 Results

### 2.1 The model

Whole-brain dynamics are modeled as the mean-field activity of neuronal populations in each brain region. We use an adapted version of the Wong-Wang model [13, 39] with a sigmoidal transfer function (equation S11). The adaption improves the biological plausibility and multistability upon the original model. Here, we briefly introduce the model; an extensive analysis of the numeric and mathematical properties of the model is provided in the Supplementary Materials (Section S4–S6 for numeric results, Section S13–S14 for analytical results). Each model region contains a pair of excitatory (E) and inhibitory (I) populations, whose activity is described by the *local model* (Figure 1a, left box; equation 1–3) in terms of the state variables *S_E_* and *S_I_*. Physically, *S_E_* and *S_I_* are interpreted as the fraction of open synaptic channels in their respective populations, i.e. the gating variables. Through local connections (*w*’s), the excitatory population excites itself with strength *w_EE_* and the inhibitory population with strength *w_EI_*, while the inhibitory population inhibits itself with strength *w_II_* and the excitatory population with strength *w_IE_*. Local models further connect to each other through a global network (Figure 1a, dashed lines), giving rise to the *global model* (right; equation 4–6). For the global model, nodes of the large-scale network correspond to anatomical regions in the human brain based on a 66-region parcellation used in [12, 42] (Figure 1b). Edge weights of the network reflect the strength of long-range structural connectivity between the brain regions (*C_ij_* in equation 6), either estimated using structural data from the Human Connectome Project [43, 44] (Section 4.5.1) or artificially constructed for comparison. The overall strength of long-range connections in the model brain is scaled by a global coupling parameter *G* (equation 6).

**Figure 1:**
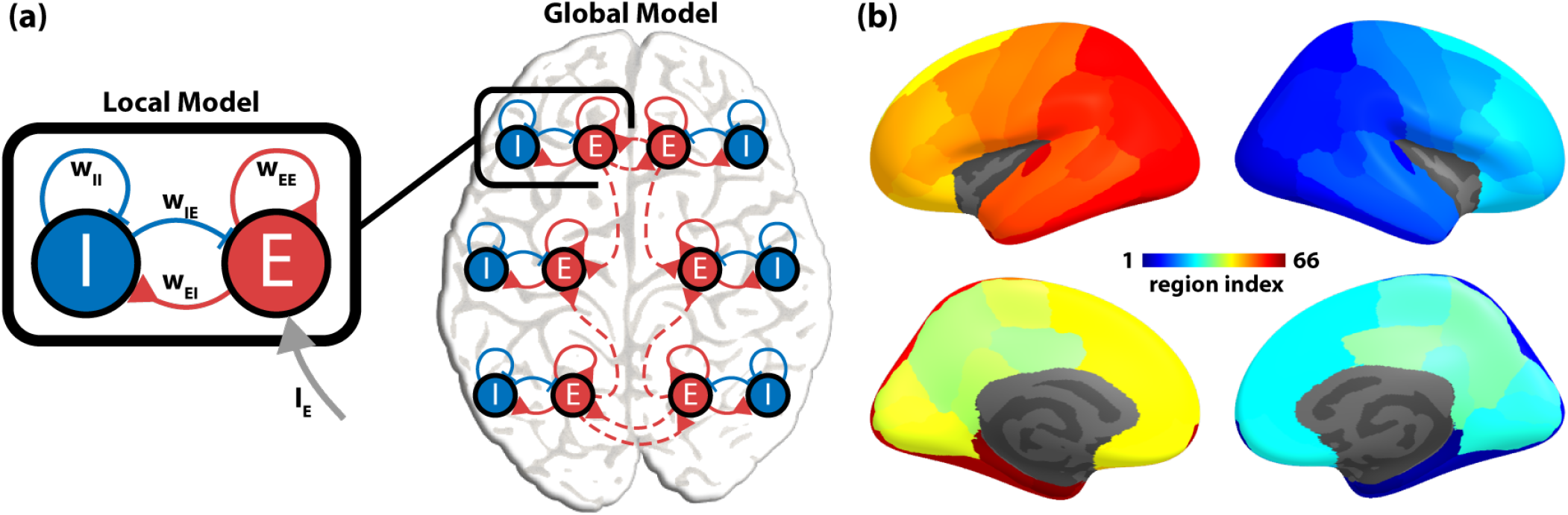
A dynamic mean-field model of the human brain. (a) The model brain (global model) consists of a network of brain regions (local model). The local model (black box) describes the interaction between two local neural populations — one excitatory (*E*) and one inhibitory (*I*). The two populations are coupled via two excitatory connections (red; *w_EE_* and *w_EI_*) and two inhibitory connections (blue; *w_II_* and *w_IE_*). The excitatory population of each brain region can further receive input (gray arrow, *I_E_*) from other regions via long-range structural connections (red dashed curves). (b) Nodes in the global model correspond to 66 anatomical regions of the human brain, which can be linked together by the human connectome (see text). Regions are indexed from 1 to 66 (1-33 on the right hemisphere, 34-66 on the left hemisphere in reverse order, following [12]). Specific region names are listed in Table S1.

In the present work, the local and global models are used in two ways: (1) to compute the repertoire of attractors using zero-finding algorithms and (2) to be numerically integrated to generate simulated brain dynamics. The former is used to characterize the overall organization of the model dynamic landscape. The latter is used to characterized local explorations of the dynamic landscape. Both aspects are compared to the human data to demonstrate the empirically relevant features. Below, we first illustrate the concept of a dynamic landscape and cross-attractor coordination using a toy example, which is followed by more realistic models to examine how structural properties across scales affect the dynamic landscape.

### 2.2 Multistable landscape of the brain shaped by structural properties across scales

Here, using a toy example (Figure 2), we first intuitively illustrate the idea of a dynamic landscape and how it may serve as a system-level description of intrinsic brain dynamics. An attractor in the toy model represents a stable pattern of activation over the whole brain, shown as boxed brains in Figure 2. The dynamic landscape determines a repertoire of attractors with possible paths of transitions between them. Such a landscape is akin to the topographical map of a land, with hills and valleys, where the attractors could be presented as valleys while the paths between the valleys can be thought of as transitions. Figure 2a represents such a toy landscape with four attractors (i-iv), each with a different whole-brain activation map. The landscape does not have to be static — as the landscape changes, some attractors (valleys) could be destroyed or created, causing a discrete change of the attractor repertoire, a.k.a. a bifurcation. Further, changes in the attractor repertoire can alter how the brain regions coordinate with each other. Two example bifurcations are shown in Figure 2b-c. In Figure 2b, only two out of four attractors are left, such that the brain can now only transition between attractor (i) and (iii), thereby leading both hemispheres to be in sync (on or off together). This coordination of brain regions (or hemispheres) can be captured by estimating the cross-attractor coordination matrix (shown in Figure 2). Similarly, in Figure 2c, three out of four attractors are left after bifurcation, leading to more complex coordination between brain regions (or hemispheres). Figure 2d-f presents a slightly more complex example, where each brain region can now take three activation values (instead of just on or off), resulting in more complex spatial activation patterns across the whole brain as well as complex coordination between brain regions. See Section 4.3 for more details about how the coordination matrix is estimated.

**Figure 2:**
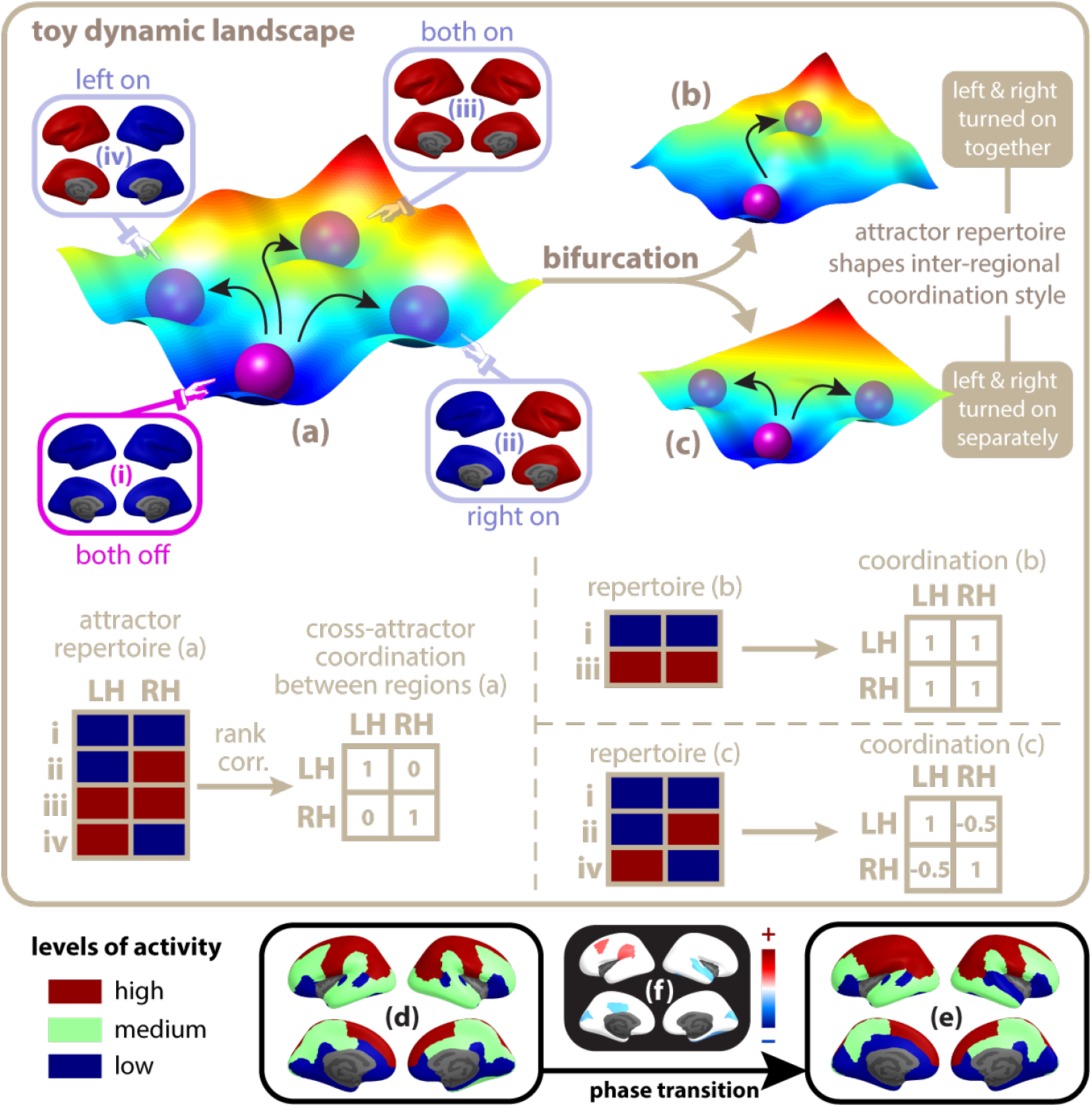
Conceptual illustration of the whole-brain dynamic landscape, bifurcation, and phase transition. A multistable dynamic landscape (a) contains multiple attractors, shown as troughs occupied by purple balls. Each attractor corresponds to a distinct pattern of activation over the whole brain (i-iv). Influenced by external input or intrinsic noise, the model brain may transition from its current state (attractor i, bright purple ball) to a different one (ii, iii, or iv, dim purple balls), indicated by black arrows. Structural features of a model brain can alter the shape of the landscape, causing some attractors to appear or disappear through a process mathematically named a bifurcation (a→b, a→c, or the reverse). By modifying the repertoire of attractors, bifurcation alters the set of possible transitions and the coordination between regions during transitions. For example, in landscape (a), the left and right hemisphere can be co-activated during a transition (i→iii), or activated separately through other transitions (i→ii, or i→iv). In contrast, in landscape (b), the left and right hemisphere can only be co-activated, and in (c), only activated separately. Numerically, a repertoire of attractors can be represented as a matrix, where each row represents an attractor and each column represents a brain region (repertoire matrix a, b, c, with entries shown as blue/red color blocks). The overall inter-regional coordination across attractors can be estimated by the rank correlation between the columns of the repertoire matrix. The resulted square coordination matrix summarizes how brain regions transition together over the entire landscape, serving as a signature of the landscape (coordination matrix a, b, c, shown to the right of each repertoire). In more complex landscapes (not shown), there are many more attractors, and they correspond to subtler patterns of activation (d,e; see also Figure 3). The coordination between brain regions during a transition is correspondingly more complex (f=e-d), with some regions co-activated (red) while others co-deactivated (blue).

Next, we present a set of more realistic examples to show how local as well as global structural connectivity of the brain can shape the dynamic landscape, its associated repertoire of attractors and their transitions. Here, we depict dynamic landscapes and their changes as bifurcation diagrams (Figure 3; see Section 4.2 for computational details). Figure 3 shows nine different bifurcation diagrams, across its three rows and columns. The rows correspond to bifurcation diagrams from: (first row: a-c) a single brain region (local model); (second row: d-f) the entire brain with uniform connectivity across all brain regions; and (third row: g-i) the entire brain with realistic connectivity across all brain regions. Thus, the second and third rows of Figure 3 aims to depict the effect of changes in global structural connectivity on the dynamical landscape. The columns, on the other hand, in Figure 3 aims to depict the effect of changes in local connectivity, i.e., level of excitation within the individual brain regions, on the dynamic landscape.

**Figure 3:**
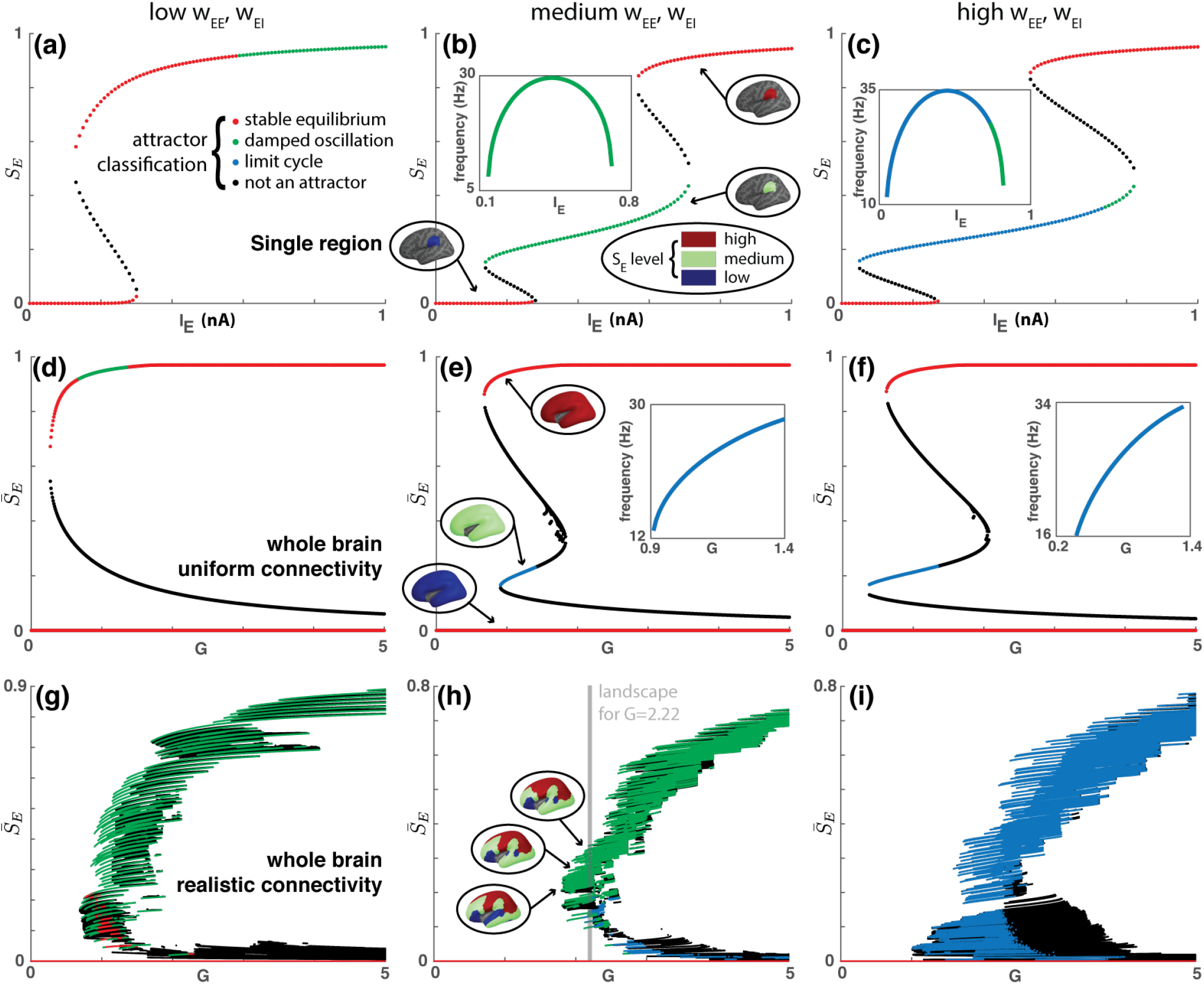
Local and global structural properties jointly determine the complexity of whole-brain dynamics. (a-c) show the bifurcation diagrams of the local model for three different types of local excitatory connectivity: (a) *w_EE_* = 0.7 and *w_EI_* = 0.35; (b) *w_EE_* = 2 and *w_EI_* = 1; (c) *w_EE_* = 2.8 and *w_EI_* = 1 (representatives of distinct dynamic regimes of the local model, Figure S1). Overall, local connectivity increases from (a) to (c). The activity of the excitatory population *S_E_* is used as an order parameter, indicating the location of each attractor. The external input *I_E_* is used as a control parameter. Each point in the diagram indicates the location of a particular fixed point. The color denotes the type of each fixed point: non-black points represent attractors, black points unstable fixed points that are not associated with a limit cycle. Horizontal stripes indicate that the attractors are changing continuously with the control parameter for a certain range. All (a)-(c) have an upper stripe and a lower stripe. (b)-(c) have an additional stripe in the middle, where the brain region oscillates. Insets of (b) and (c) show the oscillation frequency of the brain region as a function of the input current. Each stripe corresponds to a discrete level of activation for a single brain region (circled brains in b; color indicates discrete *S_E_* levels, shown in circled legend). (d)-(f) show the corresponding bifurcation diagrams for three uniform global networks, i.e. the large-scale structural connectivity *C_ij_*’s are identical between any two brain regions (equation 6). The average activity of all excitatory populations 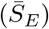 is used as an order parameter and the global coupling G (equation 6) as a control parameter. Each attractor stripe corresponds to a pattern of activation over the whole brain (circled brains in (e) show 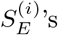 on the left hemisphere). Similarly, (g)-(i) show the corresponding bifurcation diagrams for three realistic global networks, i.e. *C_ij_*’s reflect the human connectome (see text for details). Here each vertical slice (gray line in h) contains the attractor repertoire of a fixed dynamic landscape shaped by the human connectome. Each attractor repertoire is associated with a matrix describing the coordination between brain regions across attractors (e.g. Figure 4b). See Figure 2 for a cartoon illustration of attractor repertoires and the associated cross-attractor coordination matrices.

In each bifurcation diagram, the y-coordinate of each *colored* point indicates the position of an attractor: here we use the average activity of all excitatory populations 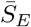 (Figure 3d-i; black points are repellers). The x-coordinate indicates the value of a control parameter, which modulates the shape of the underlying dynamic landscape: here we use the overall strength of long-range connections—the global coupling *G* (equation 6). Further, each vertical slice of a bifurcation diagram contains the repertoire of attractors and repellers in a fixed landscape (an example slice is shown in Figure 3h), corresponding to stable and unstable patterns of brain activity respectively. Lastly, an attractor traces out a horizontal “stripe” as it changes continuously with the landscape (examples shown in Figure 3d-i). A colored stripe merges with a black stripe at a bifurcation, where an attractor is annihilated by a repeller. The number of colored stripes indicates the complexity of the landscapes, i.e., more stripes indicate more attractors.

Locally wihtin each model brain region, the dynamics are controlled by local structural connectivity (*w*’s in Figure 1a; see Section S4 for detailed numeric results and Section S13 for analytical results). In particular, a single model region can switch between a rich set of dynamic regimes by varying the excitatory-to-excitatory connections (*w_EE_*) and the excitatory-to-inhibitory connections (*w_EI_*; Figure S1). Figure 3a-c show the bifurcation diagrams for three local connectivity settings from three distinct dynamic regimes (regime e, d, and a respectively in Figure S1). With an overall increase of local excitatory connectivity from (a) to (c), a single region becomes more complex—more attractors and stronger oscillatory activities.

To understand the effects of global connectivity, we first examined the brain dynamical landscape where all regions are uniformly connected with each other (Figure 3d-f). Here, stronger local excitatory connections (e,f) produce a more complex landscape (3 attractor stripes) than weak ones (d; 2 attractor stripes; see Figure S10 for even weaker local connections). These bifurcation diagrams are very similar to those of a single brain region (Figure 3a-c), in terms of the number of attractors and the presence of oscillation. In fact, the whole brain (Figure 3e) moves up and down together between discrete states of activation, very much like a single region (Figure 3b). Note that, for the global model (equation 4–6) to be multistable, a minimal amount of global coupling is required, i.e., *G* > 1 (Figure 3d-f). If the brain regions act independently (*G* = 0), both the individual regions (Figure 3a-c at *I_E_* = 0) and the whole brain (Figure 3d-f at *G* = 0) are monostable—there is only one stable pattern of activity, where the gating variables are all close to zero. This result indicates that a functionally complex brain can emerge out of the synergistic interaction between simple regions. Additional analytical and numerical results are provided in Section S14 (Multistability) for further validation and generalization.

Next, we show that in addition to the global coupling *G*, the details of inter-regional connections matter too (*C_ij_* in equation 6). Given a realistic global structural connectivity (human connectome; Figure 3g-i), the complexity of the whole-brain dynamic landscape increases dramatically: 171 attractor stripes in (g), 610 in (h), and 682 in (i) (per single-linkage clustering). Correspondingly, the patterns of activation (Figure 3h) are more complex, with greater differentiation between regions; the coordination between brain regions across attractors is consequently more flexible and subtle (as depicted in Figure 2f). The heterogeneous nature of the human connectome breaks the large-scale spatial symmetry of the model brain, creating more functional differentiation between brain regions and greater functional complexity for the whole brain. In short, the complexity of the global dynamical landscape is a joint product of strong local excitatory connection and complex topology of the large-scale network. See Section S14 for additional analytical supports.

### 2.3 Cross-attractor coordination reveals large-scale symmetry of human brain functional connectivity

In this section, using data from the Human Connectome Project (HCP) [43] we present both qualitative and quantitative results to show how cross-attractor coordination could better relate with key features of human resting functional connectivity than noise-driven within-attractor coordination. For qualitative results, we used averaged structural and functional connectivity across subjects from a smaller HCP cohort (n=11 individuals), whereas for the quantitative results, we used individual structural and functional connectivity estimates across all 100 unrelated individuals from the HCP data.

Typically, human functional connectivity (FC) is calculated from the fMRI data by estimating co-fluctuations across brain regions. The estimated resting state FC matrix usually reflects large-scale symmetry across the two hemispheres, such that brain regions across the two hemispheres highly co-fluctuate (i.e., high antidiagonal values in Figure 4a). For qualitative results, we used averaged structural and functional connectivity matrices across a small subset of the HCP n=11 unrelated individuals (see Section 4.5 and Section S9 for more details). Using averaged structural connectivity matrix, a dynamical landscape (Figure 3h) was generated and cross-attractor coordination was estimated for a chosen *G* = 2.2. At the selected G, 97 attractors were found. Mathematically, each attractor is represented by a row vector denoting the activity level of all brain regions (equation 8–9). Thus, for a chosen G, using the (#attractors × #regions) matrix we estimated the cross-attractor coordination between regions (equation 10), such that regions that co-fluctuate across attractors tend to show high cross-attractor coordination (or similarity). See Section 4.2 for mathematical details and Figure 2a-c for an intuition regarding the estimation of cross-attractor coordination matrix. As shown in Figure 4b, the dominant feature of human resting state FC, i.e., large-scale symmetry across brain regions, is well preserved in the attractor cross-coordination matrix. Such inter-hemispheric symmetry is not seen in stochastic within-attractor coordination (Figure 4d). Moreover, the pattern of within-attractor coordination is a closer reflection of the human structural connectivity (Figure 4c) than the human functional connectivity (Figure 4a).

**Figure 4:**
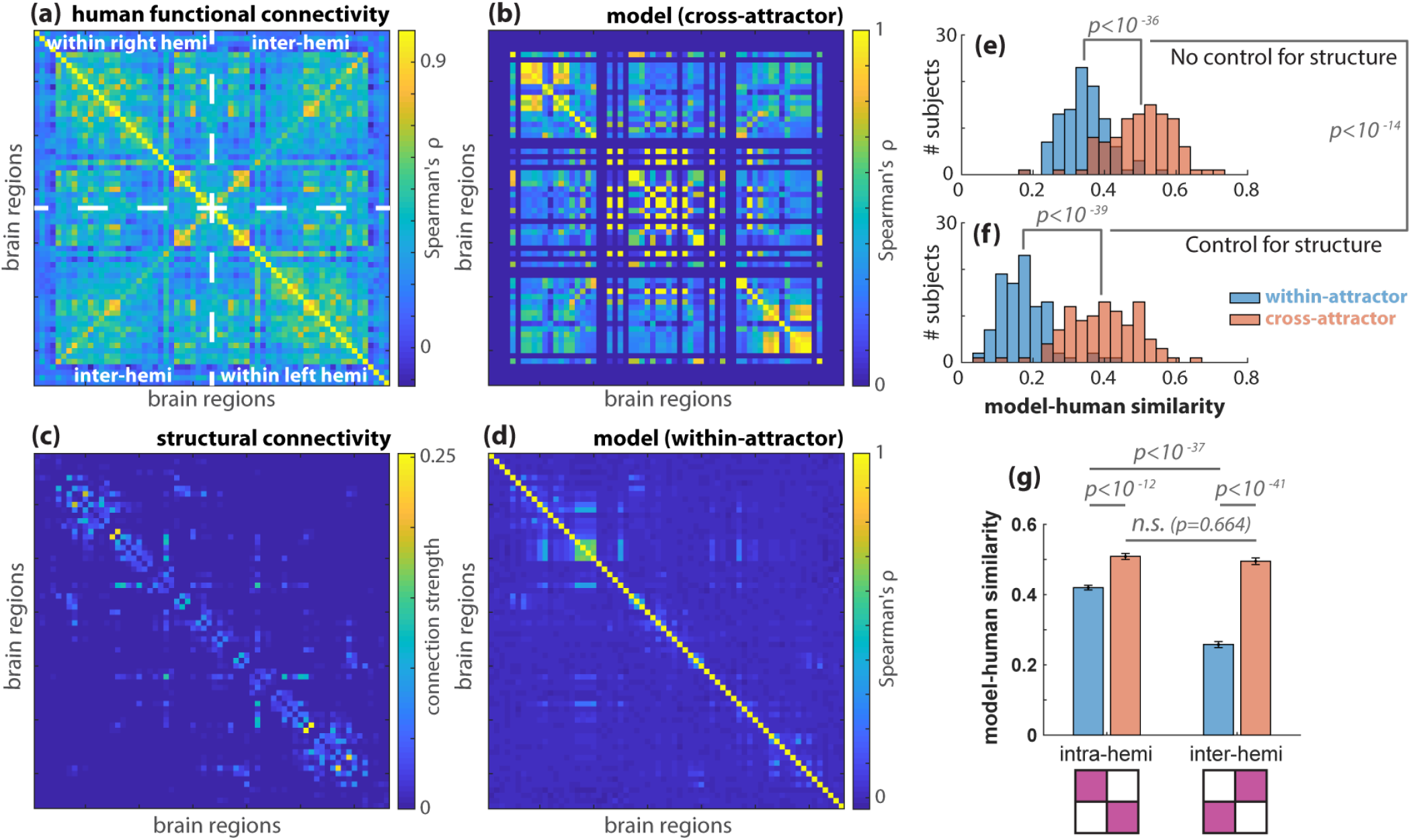
Cross-attractor coordination captures large-scale symmetry of human functional connectivity and its nonlinear dependency on structure better than within-attractor coordination. An example of human functional connectivity matrix (a) is calculated using the resting fMRI data from the Human Connectome Project [43], averaged over 11 unrelated subjects. The average structural connectivity of the same subjects is shown in (c). Regions (columns and rows) are ordered symmetrically for the left and right hemispheres (see Figure 1b) to reveal the large-scale symmetry of resting brain dynamics (ordering follows [12]; see the complete list of region names in Table S1). White dashed lines delineate the matrix (a) into four blocks, describing the functional connectivity within the right hemisphere (upper left block), within the left hemisphere (lower right), and between two hemispheres (lower left/upper right). Functional connectivity patterns within the hemispheres are similar to each other and similar to inter-hemispheric connectivity patterns. The symmetry between intra- and inter-hemispheric connectivity is well captured by inter-regional coordination in the model brain across attractors (b) (*w_EE_* = 2, *w_EI_* =1, *G* = 2.22 for local maximum model-human similarity; c.f. Figure 3h gray slice). Such symmetry is not captured by the coordination within any of the said attractors (d; best fit within-attractor coordination matrix). When examined across individuals (using n=100 unrelated HCP participants), a significantly better model-human similarity was obtained when cross-attractor coordination was used instead of within-attractor coordination (e); the difference between cross-attractor coordination and within-attractor coordination with respect to model-human similarity is even greater when partial correlation is used to control for the contribution of structural connectivity (f). Comparing intra-hemisphere and inter-hemisphere functional connectivity separately (g), we found that cross-attractor coordination captures human intra- and inter-hemisphere functional connectivity equally well (red bars), while within-attractor coordination is better at capturing intra-hemisphere than inter-hemisphere functional connectivity (blue bars). The distributions of correlation coefficients were obtained through a model fitting procedure with the same local parameters (*w_EE_* = 2, *w_EI_* = 1) while allowing the G parameter to vary from 1.7 to 3.0 by steps of 0.1.

Next, to examine the model fit on an individual basis, we ran a quantitative analysis using the n=100 unrelated-individuals cohort of the HCP data [43]. Our results confirm that cross-attractor coordination can better predict human functional connectivity than within-attractor coordination. The maximum model-human correlation for cross-attractor coordination has an average Spearman’s Rho of 0.50 (0.09), while the correlation for within-attractor coordination has an average of 0.34 (0.06) (Figure 4d). Based on within-subject paired t-test, cross-attractor coordination provided significantly better fit of FC (*t*(99) = 20.2, *p* < 10^−36^; Figure 4e). To understand how the two types of models relate to the underlying structural connectivity, we calculate the partial correlation between model coordination matrices and human functional connectivity, controlling for the linear contribution of structural connectivity. With the contribution of structural connectivity controlled, the model-human correlation for cross-attractor coordination has an average Spearman’s Rho of 0.39 (0.10), which is significantly greater than that of the within-attractor coordination 0.17 (0.07) (Figure 4f; *t*(99) = 21.9, *p* < 10^−39^). Furthermore, the difference between the partial correlation coefficients for the cross-attractor coordination and the within-attractor coordination (0.22 ± 0.10) is significantly greater than that of the regular correlation coefficients (0.16 ± 0.08) with t(99) = 9.5 and *p* < 10^−14^ (Figure 4f vs. Figure 4e), which suggests that within-attractor coordination is more linearly dependent on the structural connectivity. Finally, we show that the model-human similarity for inter-hemisphere coordination is significantly lower than that of intra-hemisphere co-ordination for within-attractor coordination but not for cross-attractor coordination (Figure 4g). In other words, unlike within-attractor coordination, cross-attractor coordination captures human intra- and inter-hemisphere functional connectivity equally well.

To understand how variability in individual model parameters translates into measures of behavior, individual parameters were correlated with a measure of fluid intelligence, an abbreviated version of the Raven’s Progressive Matrix Test (PMAT). Correlating the model parameters with measures of fluid intelligence showed one significant result. Specifically, the strength of the model-human similarity and the number of correct responses on the test was significantly correlated (Spearman’s Rho = 0.21, p = 0.04) after controlling for age and sex. All other correlations with fluid intelligence measures were not significant.

### 2.4 Structural connectivity defines the energy demands of cross-attractor coordination

In the above section, we computed the cross-attractor coordination matrices (equation 10 in Section 4.3), which only concerns whether two brain regions move up and down together across the dynamic landscape, but not how difficult or metabolically expensive such movements are. In this section, we examine the “energy gaps” between the attractors, and how they are shaped by local and structural properties of the model. Figure 5a gives a conceptual illustration of the relation between attractors and the energy gaps between them. Each attractor is associated with an average level of activity or energy (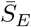; equation 11) given a fixed parameter G. There is an energy gap between each pair of adjacent attractors (equation 12; see a full technical description in Section 4.3). In the subject-average model (n=11, HCP; Figure 3g-i), the average and maximum energy gap clearly vary with local connectivity (*w_EE_*, *w_EI_*) and global connectivity (G), here summarized in Figure 5b. Quantitatively, stronger local connections reduce the energy gaps (Figure 5c,d), and stronger global connections (*G*) increase the energy gaps (Figure 5b)—local and global structural connectivity pull the energy cost in different directions.

**Figure 5:**
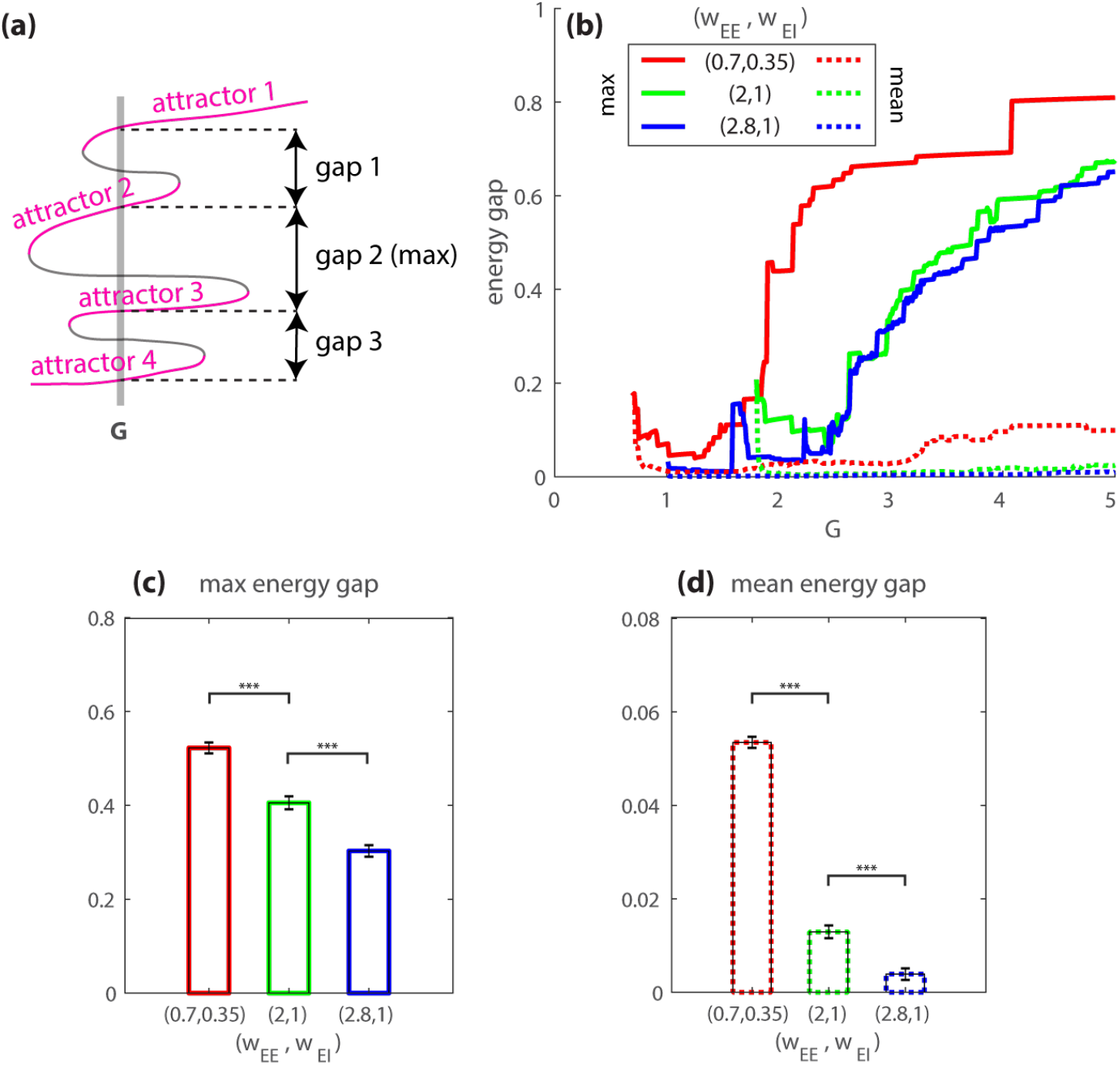
Local (*w_EE_*, *w_EI_*) and global (*G*) structural connectivity jointly shape the energy cost of cross-attractor coordination. (a) gives a conceptual illustration of the relation between attractors and energy gaps with a toy bifurcation diagram. The average and maximum gap sizes are of our interest. (b-d) summarize the size of energy gaps between attractors shown in Figure 3g-i (subject-average model with n=11). (b) Overall, the maximum (solid lines) and average energy gaps (dashed lines) increase with global coupling *G*, though there is a transient decrease when the maximum energy gap is less than 0.2. Both types of gaps decrease with increasing local connectivity *w_EE_*, *w_EI_* (c,d). (*** p<0.001 with Bonferroni correction; see Table S5–S6 for corresponding confidence intervals from the permutation tests. Error bars are standard errors.)

Next, we examine the effect of energy constraints on model fit. To incorporate the effect of energy constraints (see Section 4.3 for detailed methods), we split an ordered attractor repertoire into sub-repertoires between which the energy gap is considered too high. Thus, we obtain a sub-repertoire above the maximum energy gap, equation 13, and a sub-repertoire below the maximum energy gap, equation 14. Cross-attractor coordination matrices computed within the sub-repertoires can be considered as energy-constrained coordination patterns between brain regions. Figure 6a-c shows that such energy-constrained cross-attractor coordination (dashed lines) is more sensitive to different structural features in its ability to capture human functional connectivity. The energy constraint inflicts a greater loss of model-human similarity when the local structural connectivity is weak (area of the shaded region shrinks from Figure 6a to c, and bars in d decreases with strong local connectivity from left to right) and the global structural connectivity is strong (height of shaded regions grows with *G* in Figure 6a-c). The loss of similarity grows with the maximum gap size (Figure 6d; *ρ* = 0.96, p<10^−100^ for *w_EE_* = 0.7 and *w_EI_* = 0.35; ρ = 0.92, p<10^−100^ for *w_EE_* = 2 and *w_EI_* = 1; ρ = 0.85, p<10^−100^ for *w_EE_* = 2.8 and *w_EI_* = 1). In short, local and global structural connectivity both influence the energy costs associated with cross-attractor coordination, and thereby the model-human similarity under energy constraints. When cross-attractor coordination matrices were fitted to n=100 unrelated HCP subjects (Figure 4e), the optimal G is 2.5 ± 0.28 (Figure 6e). The corresponding maximum energy gaps average to 0.12 ± 0.08 (Figure 6f), and the corresponding mean energy gaps average to 0.009 ± 0.015 (Figure 6f). Thus, the optimal cross-attractor models in the present study are located in the regime least affected by energy constraints (Figure 6d, G<0.2).

**Figure 6:**
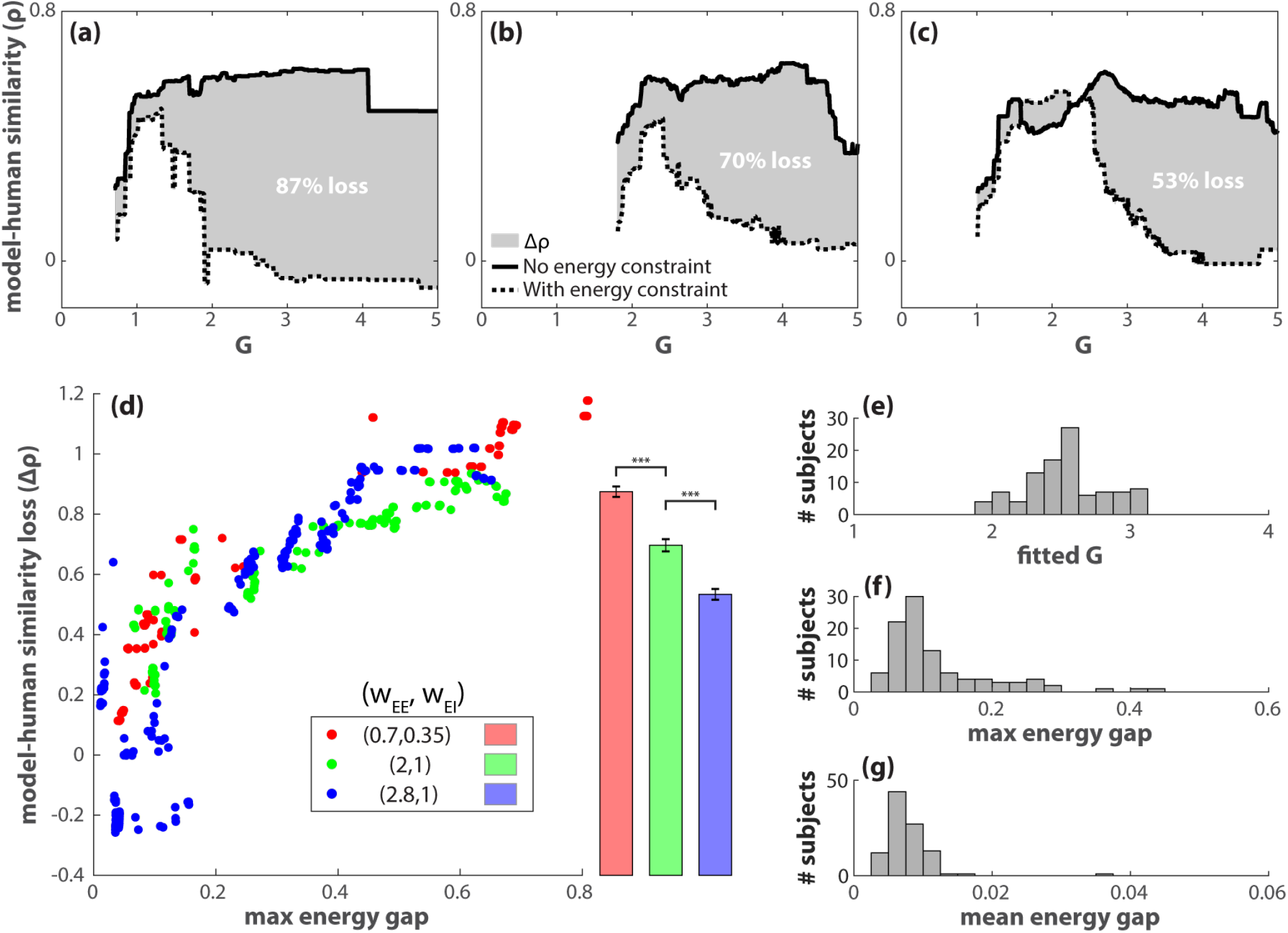
The loss of model-human similarity due to energy constraints depends on local and global structural features. (a-c) The model-human similarity for cross-attractor coordination (no energy constraint, black solid lines) is stable with respect to varying global coupling G and local excitatory connectivity *w_EE_* and *w_EI_* (a: *w_EE_* = 0.7 and *w_EI_* = 0.35, b: *w_EE_* = 2 and *w_EI_* = 1, c: *w_EE_* = 2.8 and *w_EI_* = 1). Dashed lines indicate the model-human similarity when the model is energy-constrained, i.e., it does not cross the maximum energy gap between attractors. This “energy-constrained” similarity is computed by splitting the attractor repertoire into two subrepertoires, one above the maximum energy gap and one below it (c.f. Figure 5a). The cross-attractor coordination within each subrepertoire is compared to the human functional connectivity through Spearman’s correlation. The dashed lines indicate the greater correlation coefficient (*ρ*) among the two subrepertoires. The shaded area (Δ*ρ*) indicates the loss of model-human similarity due to the energy constraint. (d) Each point in the scatter plot represents the percent loss of model-human similarity for not crossing the maximum energy gap, given a specific combination of global coupling *G* and local connectivity *w_EE_* and *w_EI_*. Overall, the loss increases with the maximum gap size, which in turn depends on G (Figure 5b). The average loss (bars in d) decreases with increasing local excitatory connectivity (*w_EE_*, *w_EI_*). When the cross-attractor model was fitted to individual subjects (using n=100 unrelated HCP participants; c.f. Figure 4e), the optimal global connectivity *G* (e) and the corresponding maximum energy gap (f) and mean energy gap (g) are low, where the cross-attractor coordination is least affected by energy constraints. (*** p<0.001 with Bonferroni correction; see Table S4 for corresponding confidence intervals from the permutation tests. Error bars are standard errors.)

## 3 Discussion

The present work examines how the brain’s multistable dynamic landscape can be shaped by structural features across scales and what features of the landscape are relevant to empirical observations. Complementing the previous stochastic noise-driven exploration approach, the present work focuses on the deterministic features of the multistable landscape and examines their empirical relevance. We demonstrate that large-scale symmetries of human functional connectivity patterns and their nonlinear dependency on the structure could be better explained by the relation between attractors in the landscape than the property of any individual attractor. Thus, the present work offers a novel cross-attractor perspective on resting brain dynamics, equipped with a computational framework to produce empirical relevant summaries of the attractor repertoire in full as well as in parts.

The functional complexity of the model brain is controlled by both local and global structural connectivity. At the level of a single isolated brain region, the dynamic repertoire can be effectively controlled by two key local structural properties: local excitatory-to-excitatory connectivity (self-excitation) and local excitatory-to-inhibitory connectivity. In the real brain, local excitatory-to-excitatory connections are particularly abundant [45], and in the model brain, they contribute indispensably to multistability (Section S13). Multistability is a key source of biological complexity from molecular to social levels [18, 19], often tied to self-excitation or positive feedback [46–48]. Manipulating the model’s local excitatory-to-excitatory connections have physical implications. For example, the manipulation can be interpreted empirically as modulating the conductance of N-methyl-D-aspartate (NMDA) receptors in local neuronal populations, using pharmacological and endogenous antagonists and agonists such as ketamine [49] and dopamine [50]. Such manipulations have been theoretically predicted and shown to affect memory capacity [51–53]. Note that the strength of local excitatory-to-excitatory connections needs to surpass a critical value to induce the transition from a monostable to a multistable regime (equation S30). In the present work, this critical value is a constant (equation S29), which depends on cellular-level properties such as the membrane time constant and the gain of the input-output response. Thus, manipulating such microscopic properties can induce, or remove, multistability from a single brain region.

At the large-scale network level, multistability can be created or amplified by the synergistic interaction between mono- or multi-stable brain regions. Different large-scale network structures have dramatically different capabilities at amplifying local complexity: a realistic global network is much more powerful than a uniform one. The human connectome breaks the spatial symmetry of the global model, whereas symmetry breaking is often a key to complex dynamics [33, 54–58]. On the other hand, the human connectome is endowed with more specific features such as modularity, small-worldness, and multiscale characteristics [1, 59–61]. A systematic study of how these features alter the geometry of the global dynamic landscape is worthy of further theoretical investigation (see Section S14).

Within the multistable landscape sculpted by the human connectome, interregional coordination across attractors exhibits key features of human functional connectivity patterns. Such cross-attractor coordination better captures human functional connectivity than within-attractor coordination—synchronization between brain regions within the same basin of attraction. This finding raises the possibility that functional connectivity patterns reflect transitions between stable brain states more than the brain states themselves. A transition-based, or cross-attractor, view on functional connectivity has several theoretical and empirical implications.

First, it provides an explanation for the large-scale symmetry of human FC, i.e., the similarity between intra- and inter-hemispheric connectivity patterns. It has been noted that within-attractor dynamics of similar models lack such symmetry, exhibiting weak inter-hemispheric FC [12, 16]. The weak inter-hemispheric FC has been attributed to an underestimation of structural connections across hemispheres using diffusion-weighted imaging. This explanation is reasonable given that within-attractor dynamics can be approximated by a linear dynamical system, which closely depends on the structural connectivity [12, 13]. Nevertheless, an alternative explanation could be that human FC implicates far-from-equilibrium dynamics where the nonlinearity cannot be ignored [15]. It is a signature of nonlinear systems that a small input does not necessarily produce a small effect. Indeed, strong functional connectivity in humans is known to exist between regions that are not directly connected [5]. As we have shown, cross-attractor coordination takes into account such nonlinear effects. Mathematically, our cross-attractor approach amounts to studying the relation between the zeros of a nonlinear function—a problem that does not admit a linear approximation. The symmetry between intra- and interhemispheric connectivity is likely to reflect a symmetry of the set of all zeros, i.e. the (roughly) invariance of the zero set under the exchange of variable indices between the homologous regions of the left and right hemispheres. The invariance of the zero set is, in turn, consequent to the invariance of the differential equations under such a left-right reflection. A full mathematical treatment of the problem is beyond the scope of the present study. Nevertheless, it invites theoretical investigations of the symmetry groups of nonlinear neural dynamical models (c.f. [55]).

The second implication is that the cross-attractor view is compatible with treating functional connectivity as both stationary and dynamical. Cross-attractor coordination, when measured over the entire dynamic landscape, is itself time-invariant. Empirically observed stability and convergence of human functional connectivity [62, 63] may reflect this invariance of the underlying landscape. The static landscape can also support dynamic functional connectivity (dFC) [15, 64]. At any given time, the possible transitions depend on the attractor currently dwelled upon. Thus, in a short time window, cross-attractor coordination is confined to a subset of attractors. In this perspective, dFC reflects the transitions between attractors in a subset of the repertoire. As a consequence, a state-based and a dFC-based representation of neural dynamics may diverge—subsets of attractors that are close in the state space may have distinct patterns of transitions, and subsets of attractors that are far apart in the state space may have similar patterns of transitions. In other words, precaution may be used when treating dFC patterns as brain states.

Finally, the cross-attractor view attaches the concept of energy costs to functional connectivity patterns. Mathematically, cross-attractor coordination matrices and energy gaps are different depictions of inter-attractor relations. Cross-attractor coordination matrices describe in which direction the attractors fall in line with each other; the energy gaps depict the spacing between attractors in a predefined direction (such a direction can represent the whole brain or a specific subnetwork). Although the pattern of cross-attractor coordination can remain similar as the landscape changes, it is not the case for its energy cost. In other words, the potential for exhibiting normal functional connectivity patterns may always be there, but the energy costs modulate the difficulty for such potential to be realized.

In summary, the present work examines intrinsic brain dynamics in terms of an underlying landscape and the repertoire of stable activity patterns it affords. Model-based analyses reveal that empirically observed functional connectivity patterns may reflect transitions between activity patterns more than the patterns themselves. The work outlines a modeling framework that emphasizes the *relation* between stable activity patterns. It is thus suitable for examining systemic changes in the brain that result in interrelated improvement or impairment in multiple cognitive and affective functions, such as in development and psychiatric disorders.

## 4 Materials and Methods

### 4.1 The present model

The local model is described by the equations,

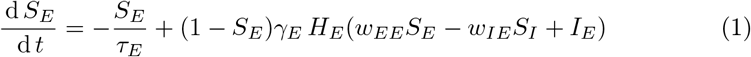

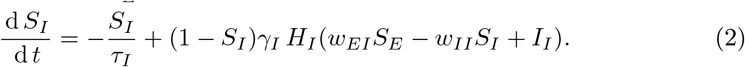

The activity of each population has a natural decay time of *τ_E_* and *τ_I_* respectively. Each population’s activity tends to increase with the fraction of closed channels (1 – *S_p_*) and the population firing rate (*H_p_*), scaled by a factor *γ_p_* for *p* ∈ {*E*, *I*}. This is described by the second term on the right-hand-side of equation 1–2. *H_E_* and *H_I_* are transfer functions that map synaptic current input to population firing rate of the excitatory and the inhibitory population respectively. In particular, they are sigmoidal functions of the form

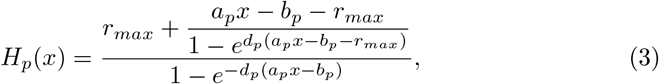

whose output increases with input monotonically and saturates at *r_max_*—the maximum firing rate limited by the absolute refractory period of neurons (around 2 ms in certain cell types [65, 66]). The specific shape of each transfer function is determined by three additional parameters *a_p_*, *b_p_* and *d_p_* (*a_p_* and *b_p_* determine the location and slope of the near-linear segment in the middle; *d_p_* determines the smoothness of the corners bordering the said near-linear segment). This transfer function is converted from Wong and Wang’s original formulation [39, 67] (a soft rectifier function, equation S6) into a sigmoidal form, while retaining the original value of parameters *a_p_*, *b_p_*, and *d_p_* (shown in Table 1). The parameters were chosen to approximate the average response of a population of spiking pyramidal cells (*p* = *E*) and interneurons (*p* = *I*) respectively, incorporating physiologically plausible parameters [39, 68]. Interaction between local populations is modulated by four coupling parameters *w_pq_* ⩾ 0 in equation 1–2, indicating the influence from the local population *p* to *q*, where *p*, *q* ∈ {*E*, *I*} (Figure 1 left box). These coupling parameters reflect the local structural connectivity. The local populations are also capable of responding to external current inputs denoted as *I_E_* and *I_I_* in equation 1–2, respectively. Importantly, such input can come from other brain regions in a globally connected network (Figure 1 right panel, dashed lines). This leads us to the global model. Formally, we substitute *I_E_* in the local model (equation 1) with a global input *I_G_* (equation 4),

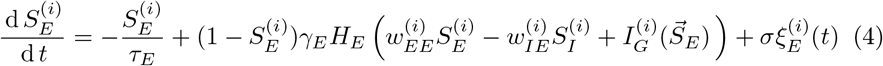

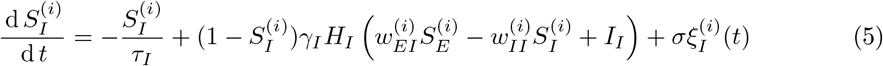

where 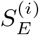 and 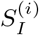 are the synaptic gating variable of the excitatory and the inhibitory population of the *i*^th^ brain region respectively, and 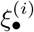 is a noise term scaled to an amplitude *σ*. The state of all excitatory populations is denoted as a vector 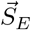, the *i*^th^ element of which is 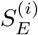. The global input to the *i*^th^ brain region depends on both its connectivity with, and the ongoing state of, other brain regions,

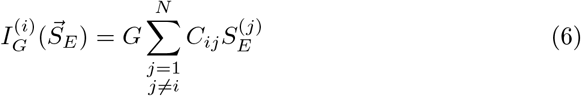

where *N* denotes the total number of brain areas, *C_ij_* ⩾ 0 the long-range structural connectivity from the *j*^th^ to the *i*^th^ brain region and *G* is a global coupling parameter that controls the overall level of interaction across brain regions. Since *C_ij_* is only intended to represent long-range connectivity, we let *C_ij_* = 0 for any *i* = *j* to preclude recurrent connections. For the effects of *G* and *C_ij_* to be independently comparable, here we impose a normalization condition on the matrix norm,

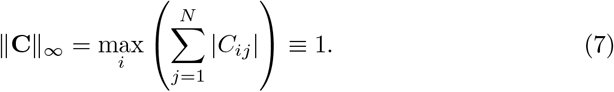

**Table 1:**
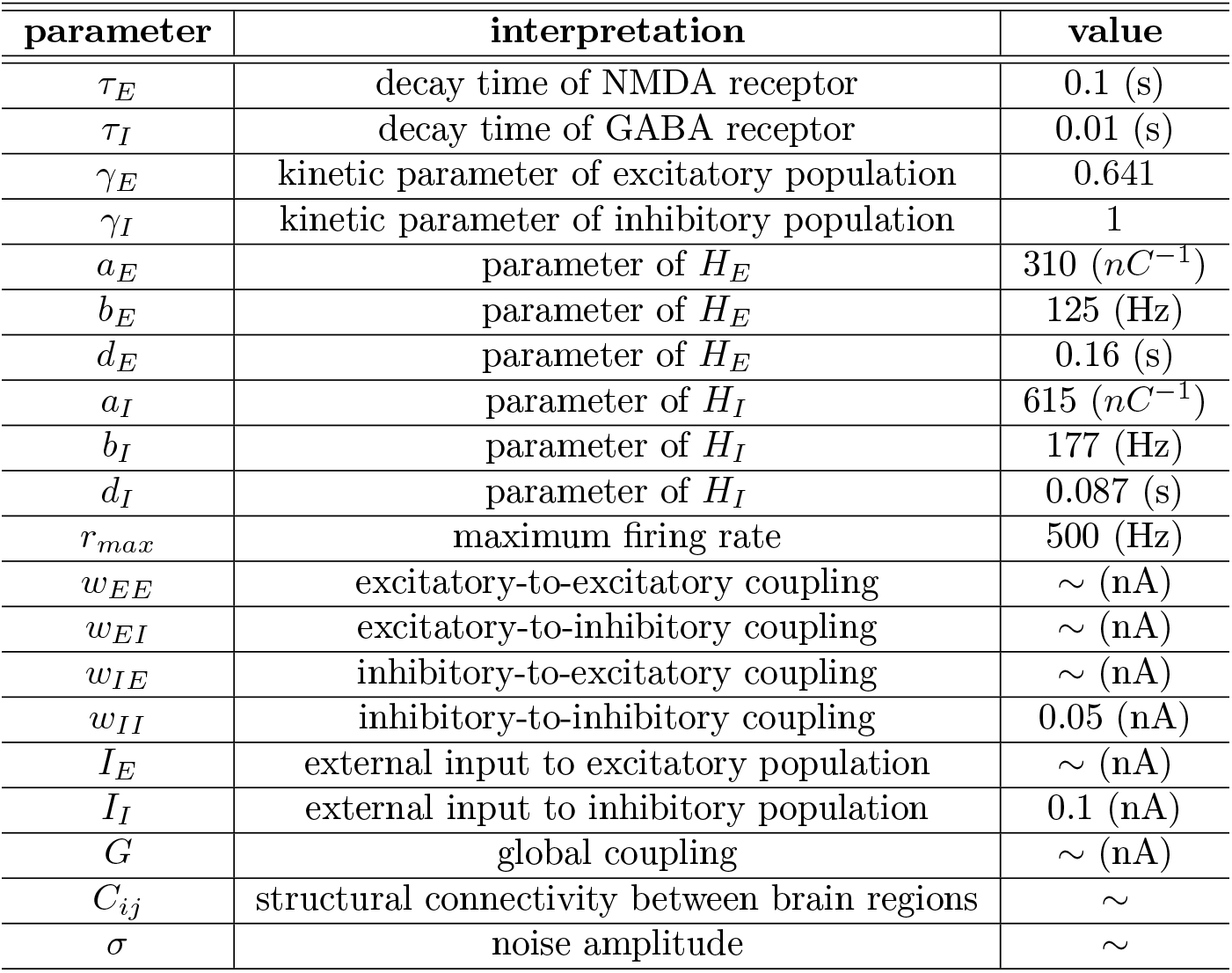
The interpretation and value of model parameters. Here we summarize the parameters used in equation 1–5. Most parameters assume a fixed value, which was introduced by [39]. A “~” indicates that this parameter is manipulated in the present study to explore the behavior of the model.

| parameter | interpretation | value |
| --- | --- | --- |
| $\tau_E$ | decay time of NMDA receptor | 0.1 (s) |
| $\tau_I$ | decay time of GABA receptor | 0.01 (s) |
| $\gamma_E$ | kinetic parameter of excitatory population | 0.641 |
| $\gamma_I$ | kinetic parameter of inhibitory population | 1 |
| $a_E$ | parameter of $H_E$ | 310 ($nC^{-1}$) |
| $b_E$ | parameter of $H_E$ | 125 (Hz) |
| $d_E$ | parameter of $H_E$ | 0.16 (s) |
| $a_I$ | parameter of $H_I$ | 615 ($nC^{-1}$) |
| $b_I$ | parameter of $H_I$ | 177 (Hz) |
| $d_I$ | parameter of $H_I$ | 0.087 (s) |
| $r_{max}$ | maximum firing rate | 500 (Hz) |
| $w_{EE}$ | excitatory-to-excitatory coupling | ∼ (nA) |
| $w_{EI}$ | excitatory-to-inhibitory coupling | ∼ (nA) |
| $w_{IE}$ | inhibitory-to-excitatory coupling | ∼ (nA) |
| $w_{II}$ | inhibitory-to-inhibitory coupling | 0.05 (nA) |
| $I_E$ | external input to excitatory population | ∼ (nA) |
| $I_I$ | external input to inhibitory population | 0.1 (nA) |
| $G$ | global coupling | ∼ (nA) |
| $C_{ij}$ | structural connectivity between brain regions | ∼ |
| $\sigma$ | noise amplitude | ∼ |

Since the global coupling parameter *G* modulates the level of input to each brain region, one would expect it to have comparable influence on the local dynamics as *I_E_* in the local model (equation 1).

### 4.2 Computation of attractors and bifurcation diagrams

The repertoire of attractors and bifurcation diagrams (Figure 3) are computed in MATLAB, utilizing the build-in function `fsolve`. Given a proper initial guess, `fsolve` finds the coordinates of a nearby fixed point of the dynamical system (e.g., the local model, equation 1–2, or global model, equation 4–5) and calculates the corresponding Jacobian matrix. Given *N* model brain regions, the spectrum 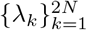 of the Jacobian matrix is used to classify the fixed points and identify which ones are attractors. The fixed point is a stable equilibrium if λ*_k_* is real and negative for all *k*. The fixed point is associated with damped oscillation if Re λ*_k_* < 0 for all *k* and Im λ*_k_* ≠ 0 for some *k*. The fixed point is associated with a limit cycle if Re λ*_k_* > 0 and Im λ*_k_* ≠ 0 for some *k* with the additional criteria that after a small perturbation from the fixed point, the time-average of the solution remains close to the fixed point. The above three types of fixed points—a stable equilibrium, a stable spiral (damped oscillation), a fixed point associated with a limit cycle (sustained oscillation)—represent attractors in the present study. All other types of fixed points are classified as unstable. For damped oscillation and limit cycles in the local model, the frequency of the oscillation (Figure S1) is defined as |Im λ*_k_*|/(2*π*).

For the local model, a 2D dynamical system, the complete characterization of all fixed points is relatively easy by searching exhaustively through a grid of initial guesses (as for Figure 3a-c). This approach becomes unfeasible when it comes to the global model due to the high dimensionality. Thus, for the global model (Figure 3d-i), we implemented a recursive search: for each value of *G*, (1) find zeros of equation 4–6 using `fsolve` given a set of initial guesses that includes, if any, the zeros for *G* – *δG* (*δG* = 0.01 for the present study) in addition to a fixed set of grid points; (2) sort the list of zeros obtained from (1) by the average of 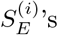 denoted as 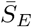; (3) use the middle points between consecutive zeros in the sorted list as initial guesses; (4) continue to use middle points between past initial guesses as new initial guesses recursively until at least one new zero is found or the recursion has reached a certain depth; (5) append the new zero(s) to the list of zeros and repeat (2)-(5) until the number of identified zeros exceeds a certain value. In the present study, we limit the maximum depth in (4) to 8 and the maximum number of zeros in (5) to 200. The set of zeros so obtained are the fixed points of the dynamical system. Each fixed point is further classified using the respective Jacobian matrix as described above to identify the attractors—a subset of the fixed points forming the attractor repertoire.

For each set of structural parameters (*G*, *C*, *w_EE_*, *w_EI_*), we represent the attractor repertoire as a M-by-N matrix,

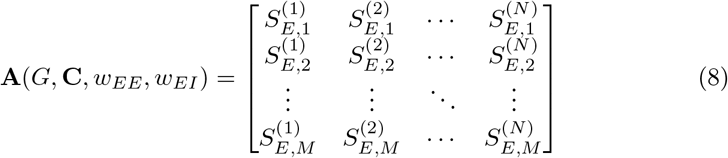

where *M* is the number of attractors, and *N* is the number of brain regions.

As parameters vary (e.g., *G* in Figure 3d-i), the attractors form discrete connected components (e.g., stripes in Figure 3) in the product of the state space and the parameter space. Attractors within the same connected component can be considered qualitatively equivalent, as they morph into each other under continuous parameter change. Understanding the relation between these connected components are critical to characterizing phase transitions and bifurcations. It is thus meaningful to define a discretized version of the attractor repertoire,

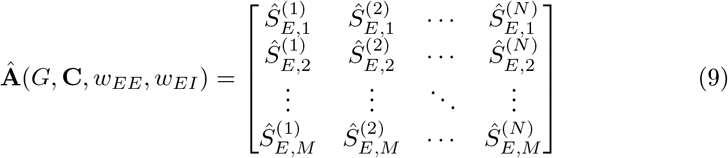

where 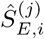 is positive integer representing a discrete level of activation which 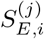 belongs to (see Figure 2 for toy examples). Each row vector in equation 9 gives the multi-index of an attractor connected component. In practice, the mapping between the continuous level 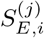 to the discrete levels of 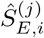 can be created by partitioning the continuous interval [0,1] at the minima of the distribution of all 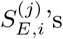 (see Section S8 for examples). Examining the properties of **A** and **Â** opens the door to systematic characterization of the underlying dynamic landscape.

### 4.3 Quantifying cross-attractor coordination and energy gaps based on the attractor repertoire

Given a discretized attractor repertoire **Â** (equation 9), the cross-attractor coordination matrix is an N-by-N matrix,

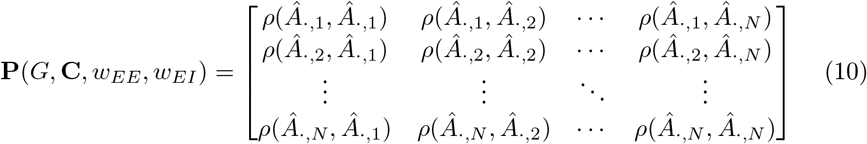

where *Â._,j_* denotes the *j*^th^ column of **Â** and *ρ*(*x*, *y*) the Spearman’s correlation between variables x and y (see Figure 2 for toy examples, Figure 4b for a more elaborate example). Spearman’s correlation is chosen to reflect the ordinal nature of the variable 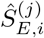. 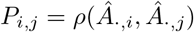 gives the level of cross-attractor coordination between model brain region *i* and *j*. The use of the *discretized* attractor repertoire **Â** ensures that coordination matrix **P** is invariant within the same dynamic regime and only changes during a bifurcation. Thus, matrix **P** connects the change of brain coordination patterns to dynamical systems concepts such as bifurcation—a qualitative change in the dynamic landscape of the model brain.

The coordination matrix **P** by itself does not explicitly concern how difficult or energy consuming these cross-attractor movements are. To incorporate energetic properties, we equip each attractor repertoire with a sequence of energy gaps. We first order the rows of the attractor repertoire matrix **A** so that the row averages descend with the row index. The row averages of the ordered repertoire matrix provide a sequence of energy levels,

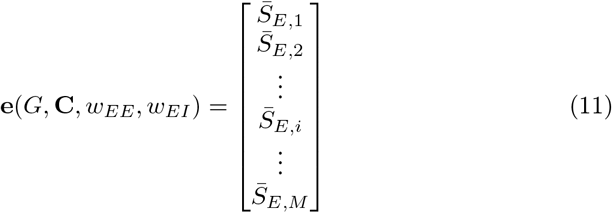

where 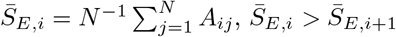 for any *i* < *M*, and *M* is the number of attractors. The corresponding energy gaps are

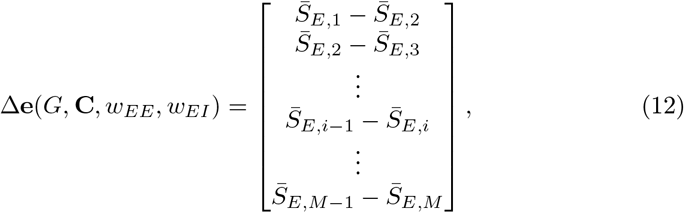

where 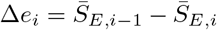 is the energy gap between the (*i* – 1)^th^ and *i*^th^ attractor in the repertoire. Physically, each energy gap Δ*e_i_* can be interpreted as the energy cost associated with keeping additional x% synaptic channels open. The sequence of energy gaps can be used to partition the attractor repertoire into submatrices. For example, if Δ*e_i_* is the maximum energy gap, one can split **A** (and its discretized version **Â**) into a repertoire above the energy gap

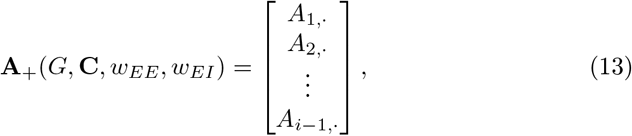

and a repertoire below the energy gap

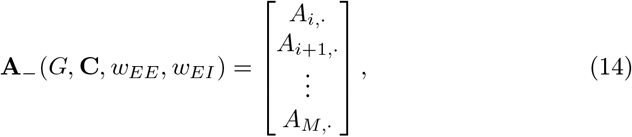

where *A_i,_*. is the *i*^th^ row of the ordered repertoire matrix **A**. Each of the subrepertoires **A**_+_ and **A**_−_ is associated with its own cross-attractor coordination matrix, say, **P**_+_ and **P**_−_ (equation 10). **P**_+_ and **P**_−_ can be considered as the “energy-constrained” coordination patterns where the model brain is not allowed to cross the energy gap Δ*e_i_*. The same line of analysis is applicable to a sub-network of the model brain (e.g., the default mode network) by constructing a reduced repertoire matrix that only contains a subset of the columns in **A**. These selected columns map to the brain regions within the sub-network. This series of analysis applied to attractor repertoire matrices and sub-matrices provides a systematic and simple way to characterize brain dynamic landscape and inter-regional coordination.

### 4.4 Estimating within-attractor coordination through simulations

Within-attractor coordination is estimated using conventional methods of numeric simulation. In the present work, for each attractor in the repertoire, the dynamical system (equation 4–6) is integrated using stochastic Heun’s method, with a time step of 1 ms, a moderate level of noise *σ* = 0.01 and a duration *T* = 864 s (14 min 33 s to match the human data [43]). The exact coordinates of the attractor are used as the initial conditions such that the simulated dynamics reflects the noise-driven exploration near that attractor. Conventional correlation analysis is then applied to the simulated time series of the excitatory populations 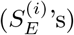 to obtain the functional connectivity matrix (e.g., Figure 4c). Spearman’s correlation is used in accordance with the computation of cross-attractor coordination matrix (**P**, equation 10).

### 4.5 Data and methods of analysis

#### 4.5.1 Human structural data

The human structural connectome used in the present study is from the S1200 Release from the Human Connectome Project (HCP) [43]. The average connectome of 11 unrelated subjects were used for the first qualitative analysis (it has been shown that averaging over 5 subjects is sufficient [12, 15]), while the individual connectome of 100 subjects obtained from a previous study [44] were used for the second quantitative analysis. For both analyses, the subject-level connectome data are based on the Desikan-Killiany parcellation [69] obtained from [44], retaining the 66 ROIs used in [42] and [12] (Figure 1b). The original diffusion imaging (dMRI) data were obtained using a customized Siemens 3T scanner at Washington University in St. Louis, with a standard 32-channel head coil, with TR = 5520 (ms), TE = 89.5 (ms), 1.25 (mm) isotropic voxels, b=1000, 2000, 3000 (s/mm^2^). T1 images were obtained using 3D magnetization-prepared rapid gradient echo sequence (MPRAGE) with TR = 2400 (ms), TE = 2.14 (ms), and 0.7 (mm) isotropic voxels. The HCP minimally processed data were further processed using *MRtrix3*, including biasfield correction, multi-shell multi-tissue constrained spherical deconvolution with a maximum spherical harmonic degree 8. 10 million probabilistic streamlines were generated for each subject using the 2^nd^-order Intergration over Fibre Orientation Distributions algorithm (iFOD2) [70] and anatomically-constrained tractography (ACT) [71] (FOD amplitude threshold = 0.06, step size = 0.625 mm). Each streamline was assigned a weight using spherical-deconvolution informed filtering of tractograms (SIFT2) [72]. Connection strengths between ROIs are summed weights of the associated streamlines. Intra-ROI connections are removed. Subjects’ connectivity matrices are normalized according to equation 7 before and after averaging.

#### 4.5.2 Human functional data

Human functional connectivity used in the present study is estimated using the resting-state fMRI (rsfMRI) data of the same subjects from the Human Connectome Project [43] as aforementioned ones from the structural connectivity. rsfMRI scans were acquired using EPI sequences with TR = 720 (ms), TE = 33.1 (ms), flip angle = 52°, voxel size = 2.0 (mm, isotropic), multiband factor = 8. Four runs of rsfMRI scan were obtained from each subject in 2 separate days (2 runs in each day with opposite phase-encoding direction: RL and LR). Each run last 14 min 33 s (1200 TR).

For the first analysis, unprocessed data were downloaded from the Human Connectome Project database (https://db.humanconnectome.org) and preprocessed using *fMRIPrep* 1.4.0 ([73]; [74]; RRID:SCR_016216), which is based on *Nipype* 1.2.0 ([75]; [76]; RRID:SCR_002502). First, a reference volume and its skull-stripped version were generated using a custom methodology of *fMRIPrep*. A deformation field to correct for susceptibility distortions was estimated based on two echo-planar imaging (EPI) references with opposing phase-encoding directions, using `3dQwarp` [77] (AFNI 20160207). Based on the estimated susceptibility distortion, an unwarped BOLD reference was calculated for a more accurate co-registration with the anatomical reference. The BOLD reference was then co-registered to the T1w reference using `bbregister` (FreeSurfer) which implements boundary-based registration [78]. Co-registration was configured with nine degrees of freedom to account for distortions remaining in the BOLD reference. Head-motion parameters with respect to the BOLD reference (transformation matrices, and six corresponding rotation and translation parameters) are estimated before any spatiotemporal filtering using `mcflirt` [FSL 5.0.9, 79]. The BOLD time-series were resampled to the *fsaverage* surface space. Several confounding time-series were calculated including framewise displacement (FD), DVARS and three region-wise global signals. FD and DVARS were calculated for each functional run, both using their implementations in *Nipype* [following the definitions by 80]. The three global signals were extracted within the CSF, the WM, and the whole-brain masks.

Nuance regressions were performed on detrended, preprocessed BOLD time series in the *fsaverage* space (FreeSurfer), following procedures in [80]. Regressors include 6 motion parameters, CSF signal, WM signal, and their first derivative and second power. Frames with FD>0.2 mm are censored. Spline-interpolated signals are band-pass filtered between 0.009 and 0.08 Hz, and averaged within ROIs based on the Desikan-Killiany parcellation [69]. 66 Regions in [42] are retained and ordered according to [12] (Figure 1b). Functional connectivity between ROIs are estimated using Spearman correlation between z-scored time series for each rsfMRI run of each subject. The connectivity matrices are then averaged across all runs/subjects in Day 1 and in Day 2 separately. The average functional connectivity matrix from Day 1 is used in all comparisons with the model. The average functional connectivity matrix from Day 2 is used to assess the reliability of the estimation.

For the second analysis, an alternative preprocessing pipeline that is standard for individual rsfMRI preprocessing was applied. Specifically, minimally preprocessed rsfMRI volumetric data from the HCP database was downloaded and further denoised through application of a published spatial independent component rejection procedure [81–83]. The resulting rsfMRI timeseries was then converted to ROI space by averaging voxels part of the Desikan-Killiany atlas in volume space [84]. Relative to the list in Table S1, there were six missing ROIs, which are the left and right temporal pole, the left and right frontal pole, and the left and right bank of the superior temporal sulcus. Similar to the case with the 11 subjects, there were 4 runs per subject (2 runs with opposite encoding for 2 separate days). To calculate the functional connectivity, ROI time series were z-scored and then correlated with Spearman correlation between each pair of ROIs. The resulting 4 connectivity matrices were averaged to obtain one final connectivity matrix per subject.

#### 4.5.3 Individual subject simulation and model fitting

To find the optimal human-model fitting on an individual basis, simulations were run with one representative set of local parameters (*W_EE_* = 2, *W_EI_* = 1), while the global coupling parameter was varied from 1.7 to 3.0 with step sizes of 0.1. This set of parameter configurations were chosen with considerations for the run time while capturing the individual variability for the optimal value of G. Additional details of the simulation, including computation of the within and cross attractor coordination, and energy gaps between attractors, were identical to the analysis of the 11 subject averaged data.

Spearman’s correlation was used to quantify the model-human correlation. The optimal G parameter for each individual was determined based on the maximum correlation for the cross-attractor coordination. Within-attractor coordination matrices were determined for individual attractors at that optimal G and used to find the highest model-human correlation for within-attractor coordination. To compare the model fit between within and cross-attractor coordination on an individual level, a paired t-test was applied between the model-human correlation for the two across the 100 individuals.

#### 4.5.4 Correlating with measures of fluid intelligence

An abbreviated 24-items version of the Raven’s Progressive Matrix Test Form A was used for evaluating fluid intelligence Duncan, 2000; Bilker, 2012 and scored with the number of correct responses (PMAT24_A_CR). Spearman’s correlation was calculated between model parameters and PMAT24_A_CR while controlling for age and sex. Model parameters of interest include the maximum correlation value, the associated optimal G parameter, and the maximum energy gap at the optimal G.

## 5 Acknowledgements

This work is supported by a NIH Director’s New Innovator Award to M.S. (MH119735).

## Supplementary Materials

### S1 Region indices and names

**Table S1:**
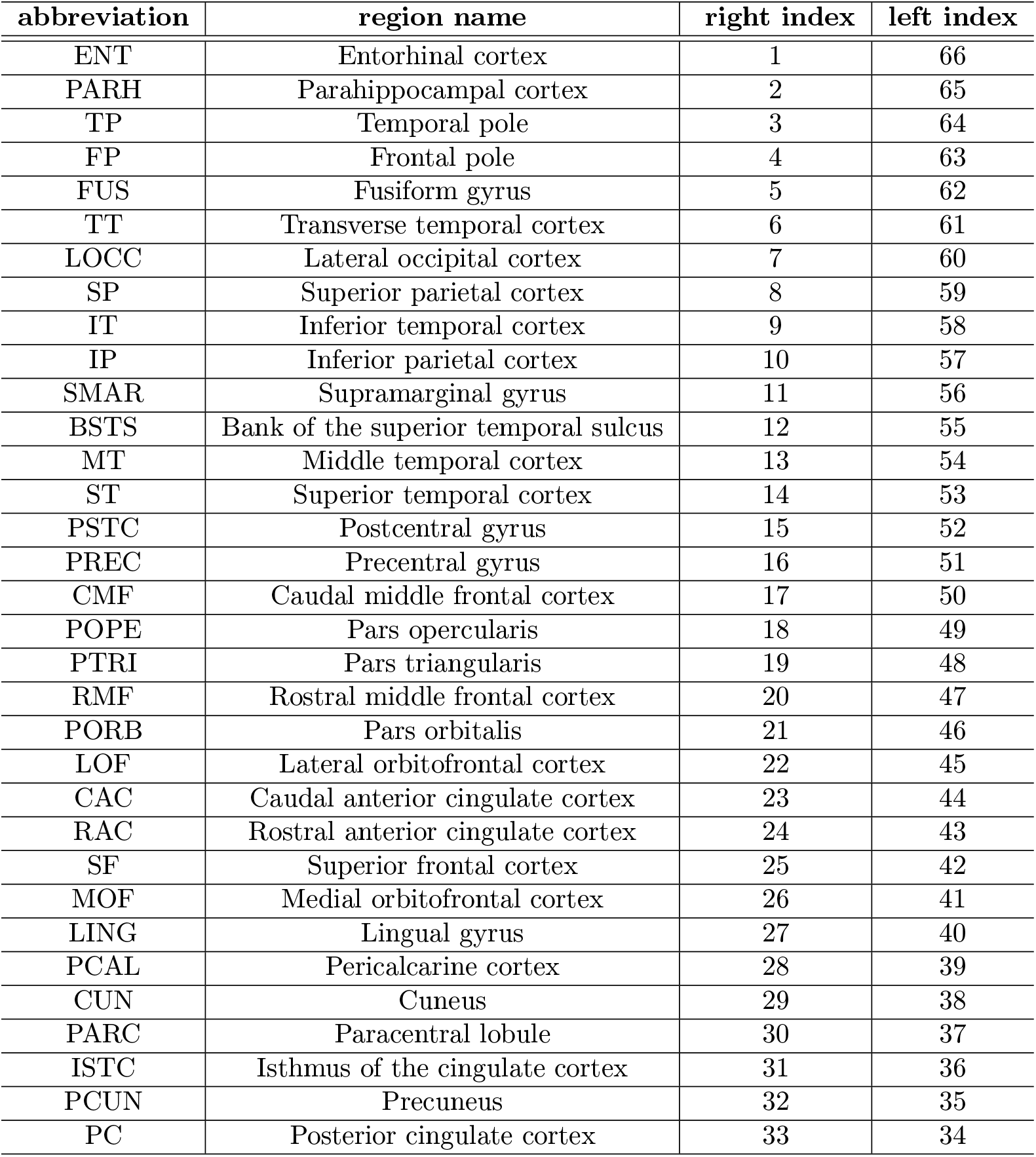
Names and indices of brain regions. As in [1], the brain is parcellated into 66 regions as shown in Figure 1b. Here is a list of the specific region names corresponding to each region index. Region 1-33 are on the right hemisphere, and region 66-34 are the homologous regions on the the left hemisphere.

| abbreviation | region name | right index | left index |
| --- | --- | --- | --- |
| ENT | Entorhinal cortex | 1 | 66 |
| PARH | Parahippocampal cortex | 2 | 65 |
| TP | Temporal pole | 3 | 64 |
| FP | Frontal pole | 4 | 63 |
| FUS | Fusiform gyrus | 5 | 62 |
| TT | Transverse temporal cortex | 6 | 61 |
| LOCC | Lateral occipital cortex | 7 | 60 |
| SP | Superior parietal cortex | 8 | 59 |
| IT | Inferior temporal cortex | 9 | 58 |
| IP | Inferior parietal cortex | 10 | 57 |
| SMAR | Supramarginal gyrus | 11 | 56 |
| BSTS | Bank of the superior temporal sulcus | 12 | 55 |
| MT | Middle temporal cortex | 13 | 54 |
| ST | Superior temporal cortex | 14 | 53 |
| PSTC | Postcentral gyrus | 15 | 52 |
| PREC | Precentral gyrus | 16 | 51 |
| CMF | Caudal middle frontal cortex | 17 | 50 |
| POPE | Pars opercularis | 18 | 49 |
| PTRI | Pars triangularis | 19 | 48 |
| RMF | Rostral middle frontal cortex | 20 | 47 |
| PORB | Pars orbitalis | 21 | 46 |
| LOF | Lateral orbitofrontal cortex | 22 | 45 |
| CAC | Caudal anterior cingulate cortex | 23 | 44 |
| RAC | Rostral anterior cingulate cortex | 24 | 43 |
| SF | Superior frontal cortex | 25 | 42 |
| MOF | Medial orbitofrontal cortex | 26 | 41 |
| LING | Lingual gyrus | 27 | 40 |
| PCAL | Pericalcarine cortex | 28 | 39 |
| CUN | Cuneus | 29 | 38 |
| PARC | Paracentral lobule | 30 | 37 |
| ISTC | Isthmus of the cingulate cortex | 31 | 36 |
| PCUN | Precuneus | 32 | 35 |
| PC | Posterior cingulate cortex | 33 | 34 |

### S2 Relation to the Wilson-Cowan model

Formally, the above model can be considered a special variant of the Wilson-Cowan model [2, 3]. Though the specific interpretation of certain parameters differ, the two models describe similar dynamic mechanisms of population-level interaction. The Wilson-Cowan model, in its initial form [2], concerns the dynamics of a pair of interacting excitatory and inhibitory neuronal populations. The activities of the two populations are denoted as *E*(*t*) and *I*(*t*)—the proportion of firing excitatory/inhibitory cells averaged over a period of time (the refectory period). The model takes the form

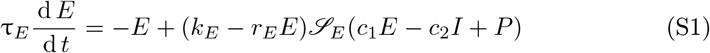

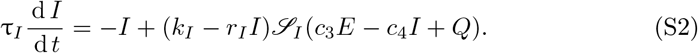

*τ_E_* and *τ_I_* are time constants of the dynamics of the excitatory and inhibitory population respectively. *c*_•_’s are the coupling parameters between the two population. Coefficients *k*_•_, and *r*_•_ result from a temporal coarse-graining procedure in the initial derivation (see [2] for detail). 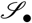 is a sigmoidal transfer function, rising monotonically from 0 to 1 with non-negative input. *P* and *Q* are external inputs to their respective populations. If we divide both sides of equation S1–S2 by the time constants, we are looking at the same general form as equation 1–2.

The main difference is between the respective transfer functions. Wilson and Cowan [2] chose a particular form of 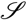 for mathematical analysis:

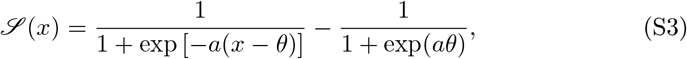

where parameter *a* determines the maximum slope of the function 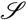 and parameter *θ* the location of the maximum slope. Technically, Wilson and Cowan [2] only requires 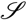 to be of a general sigmoidal form. It may reflect the average response of a population of neurons with heterogeneous firing thresholds or heterogeneous afferent connections. The distribution of the said thresholds or connections is reflected in the parameters *a* and *θ*.

In other words, the choice of the transfer function and the parameters is *non-specific* to a predefined microscopic model. Moreover, Wilson and Cowan [2] took a function-oriented approach to analyzing the model. The key was whether the model was able to produce fundamental behaviors expected from a neural model—multistability, hysteresis, and oscillation—for *some* specific choice of parameters and transfer function. Qualitative conclusions from their analysis depend on the general geometric properties of the transfer function rather than the specific form of equation S3.

The transfer function of the present model (equation 3) follows the general geometric properties assumed by Wilson and Cowan [2]. The difference is that the parameters in equation 3 are associated specifically with a microscopic model [4], a network of leaky integrate-and-fire neurons with biologically plausible parameters, as inherited from the reduced Wong-Wang model [5, 6]. This choice provides a channel of correspondence between parameters of the models at different scales of description. To expand on this point, we next elaborate on the connection between the present model and the reduced Wong-Wang model.

### S3 Relation to the reduced Wong-Wang model

The present model is also a variant of the Wong-Wang model [5] and its highdimensional generalizations, here referred to as the reduced Wong-Wang model [1, 6, 7]. In particular, we consider the model of whole-brain dynamics [6, 7],

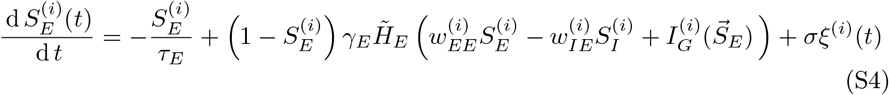

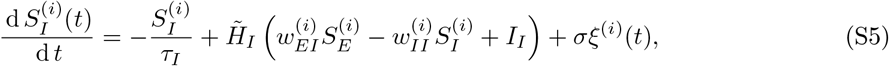

following the same notations as in equation 4–6, where

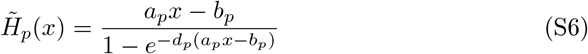

with *p* ∈ {*E*, *I*} denoting the excitatory and the inhibitory population respectively. The parameters *a_p_*, *b_p_* and *d_p_* were chosen such that 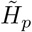 approximates the average firing rate of an ensemble of leaky integrate-and-fire neurons receiving uncorrelated noisy inputs.

More specifically, the sub-threshold dynamics of the membrane potential *V*(*t*) of each neuron can be described as

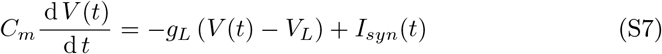

where *C_m_* is the membrane capacitance, *g_L_* the leak conductance, and *V_L_* the resting potential of the membrane. The total synaptic input current *I_syn_*(*t*) is a random process with an average and standard deviation *σ_C_*. When *V*(*t*) reaches a threshold *V_th_*, the neuron emits a spike after which the membrane potential returns to a reset voltage *V_reset_* and stays there for a duration *τ_ref_*, i.e. the refractory period.

The average firing rate *ν* of an ensemble of such neurons can be derived from the Fokker-Planck approximation that describes the evolution of the membrane voltage distribution of an ensemble of neurons (see e.g. [8, Section 1], [9] for descriptions of the Fokker-Planck approach). This eventually leads to the first-passage time equation (average time for crossing the threshold),

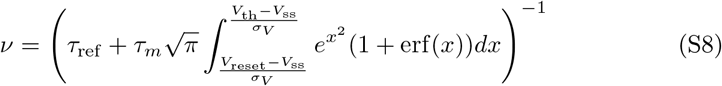

where *τ_m_* = *C_m_*/*g_L_* is the membrane time constant, 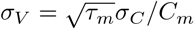 the standard deviation of the depolarization, erf(*x*) the error function

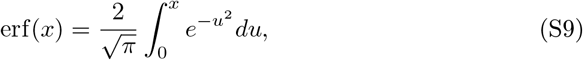

and *V_ss_* the steady state voltage

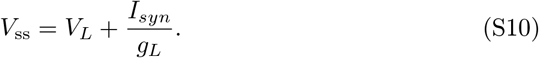

The transfer function employed by Wong and Wang [5, 10], i.e. equation S6 with appropriate choice of parameters, is a good approximation of equation S8 when the input level is low.

Thus, the first passage equation S8 provides a bridge between the transfer function (equation S6) and the single-cell level model (equation S7) incorporating realistic biophysical parameters (Table S2). In other words, it allows one to use empirically measurable quantities at the neuronal level to directly constrain the the transfer function and the entire model. This is a major difference with the Wilson-Cowan model [2, 3].

**Table S2:** Biophysical parameters of a single leaky-integrate-and-fire neuron. If two parameter values are provided in the right column, the first value is for a generic pyramidal cell and the second is for a generic interneuron. Differences between the biophysical parameters of different cell types lead to differences in the transfer functions (equation S6).

| parameter | interpretation | value |
| --- | --- | --- |
| $C_m$ | membrane capacitance | 0.5, 0.2 (nF) |
| $g_L$ | leak conductance | 25, 20 (nS) |
| $\tau_m$ | membrane time constant | 20, 10 (ms) |
| $\tau_{ref}$ | refractory period | 2 (ms) |
| $V_L$ | resting membrane potential | -70 (mV) |
| $V_{th}$ | threshold for firing | -50 (mV) |
| $V_{reset}$ | reset potential | -55 (mV) |

According to the first passage equation S8, the firing rate *ν* is a sigmoidal function of the input, which saturates at *r_max_* ≡ 1/*τ_ref_*. This is not the case, however, for the transfer function 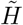 (equation S6).To make the transfer function a better approximation of the first passage equation and at the same time retain the mapping between their parameters, we can simply convert the transfer function by substituting the numerator of 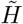 as below

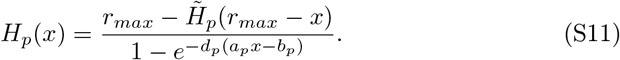

Thus, we obtain the transfer function used in the present model (equation 3). *H_p_* matches 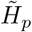 for low levels of input but flattens out eventually at *r_max_* as one would expect from equation S8.

In short, the present model is endowed with the geometric properties of the Wilson-Cowan model [2, 3] and at the same time consistent with the neuronal level-to-population level mapping of the reduced Wong-Wang model [5, 6].

### S4 Local structural connectivity controls dynamic repertoire of an isolated brain region

**Figure S1:**
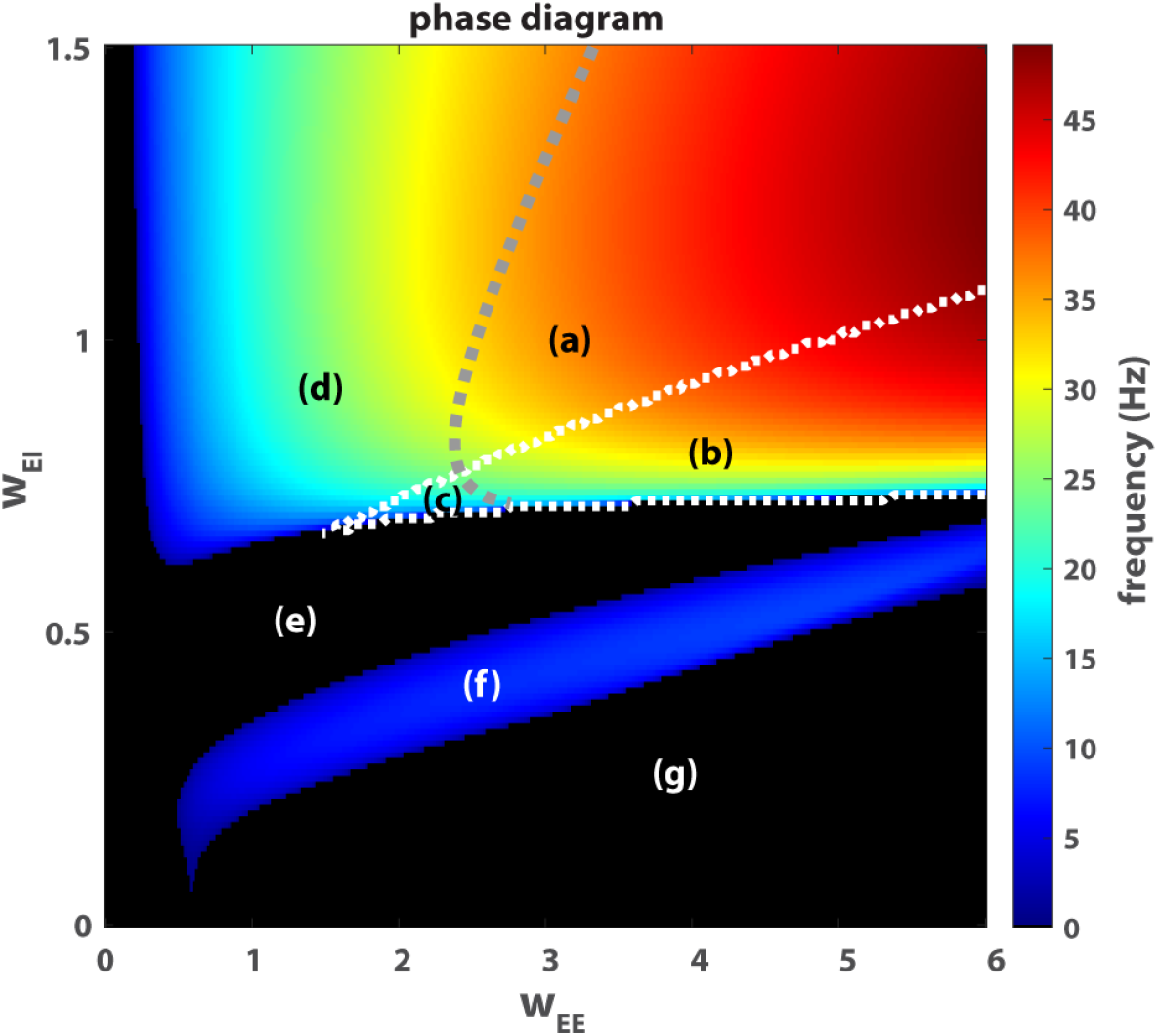
Local dynamics controlled by the strength of excitatory-to-excitatory connection *w_EE_* and excitatory-to-inhibitory connection *w_EI_*. (a)-(g) are seven different dynamic regimes of the local model (equation 1–2) in the 2-dimensional parameters space (*w_EE_*, *w_EI_*). Here the local inhibitory-to-excitatory connectivity *w_IE_*—the inhibitory feedback—is matched to the excitatory-to-excitatory connectivity, i.e. *w_IE_* = *w_EE_*; and *I_E_* = 0.382 as in [6]. Black areas (e, g) are the regimes of stable equilibrium. Colored areas are the oscillatory regimes: (a)-(b) for limit cycles and (c), (d), (f) for damped oscillations. The color reflects the frequency of oscillation. A gray dashed line indicates the Hopf bifurcation. The triangular area enclosed by white dashed lines (saddle-node bifurcation) is the bi-stable regime (b, c). A typical phase portrait from each regime is provided in Figure S2.

The local model exhibits a rich repertoire of dynamical features, including multistability (b, c in Figure S1, S2), damped oscillation (c, d, f), and limit cycles (sustained oscillation; a, b). Mathematical analysis of the local model (Section S13 in Supplementary Materials) shows that nonlinearity in the dynamics can essentially be controlled by two local structural properties: the strength of excitatory-to-excitatory connection *w_EE_* and the strength of the excitatory-to-inhibitory connection *w_EI_*. Geometrically, the two structural properties “twist” the nullclines (dashed lines in Figure S2). Specifically, stronger *w_EE_* introduces a deeper twist and fold of the red nullcline (compare Figure S2a and d), whereas stronger *w_EI_* introduces a more vertical twist of the blue nullcline (compare Figure S2d and e). These twists are the key sources of dynamic complexity—multistability and oscillation. For example, when *w_EE_* is sufficiently large (equation S30), multistability becomes possible: the folded red nullcline allows for multiple intersections with the blue nullcline, and potentially a greater number of attractors (compare Figure S2c to a; see analytical results in Section S13: Multistability). When *w_EI_* is sufficiently large (equation S38), oscillatory activity becomes possible (analytical results in Section S13: Oscillation). Moreover, the combination of large *w_EE_* and *w_EI_* gives rise to sustained oscillation (equation S61). The characteristic frequency of such oscillation further depends on the specific values of *w_EE_* and *w_EI_*. Note that the general qualitative effects of these two local structural properties are consistent with those of the Wilson-Cowan model, but the specific boundaries at which transitions occur are determined by the biophysical constraints inherited from the reduced Wong-Wang model (see equations S38, S61). Analytical results (Section S13) provide detailed quantification of how these boundaries are shifted by different local structural properties.

To maintain a sufficient twist in the red nullcline (red dashed line in Figure S2) and associated multistability and oscillation, inhibitory-to-excitatory feedback *w_IE_* needs to be proportional to self-excitation *w_EE_* (c.f. equation S20). In the present study, we simply let *w_IE_* = *w_EE_*. This equality is a simpler alternative to the Feedback Inhibition Control adopted in [6] in both numerical and mathematical analyses.

**Figure S2:**
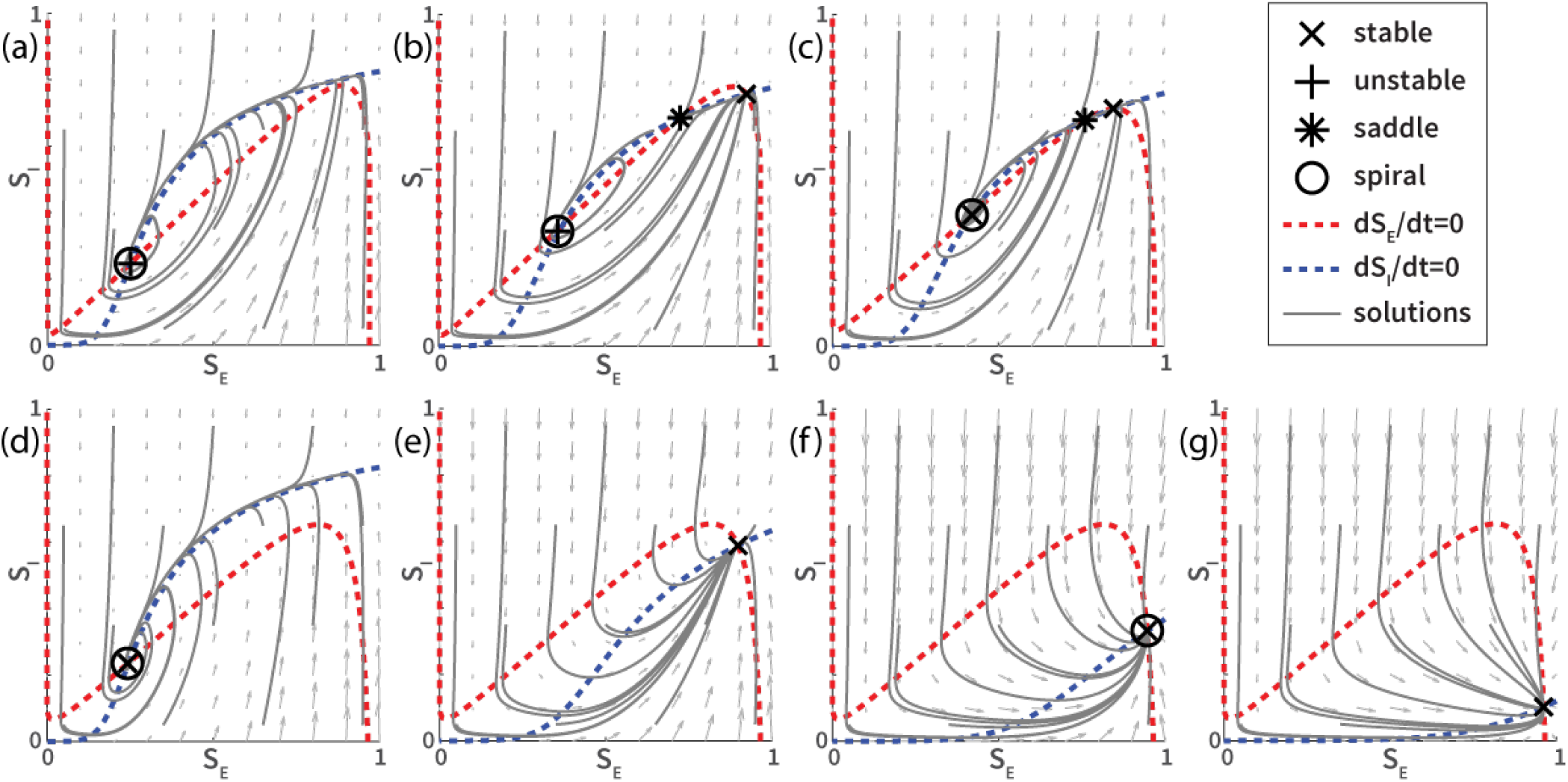
Example phase portraits from different regimes of the local model. Phase portraits (a) to (g) are examples chosen from the corresponding regimes in Figure S1. The specific parameters defining local structural connectivity are (a) *w_EE_* = 4, *w_EI_* = 1; (b) *w_EE_* = 4, *w_EI_* = 0.8; (c) *w_EE_* = 2.3, *w_EI_* = 0.75; (d) *w_EE_* = 1.5, *w_EI_* = 1; (e) *w_EE_* = 1.5, *w_EI_* = 0.5; (f) *w_EE_* = 1.5, *w_EI_* = 0.3; (g) w_EE_ = 1.5, *w_EI_* = 0.2. The vector fields (arrows) reflect the underlying dynamics at different points in the state space. Gray trajectories following the vector fields are solutions of the local model (equation 1, 2) given a fixed sets of ten different initial conditions. Nullclines (dashed lines) indicate where the flow of the dynamics is either purely vertical (red) or purely horizontal (blue). The intersections between the nullclines are the fixed points. Different types of fixed points are labeled with different markers (see legend). A fixed point is stable (×) if nearby trajectories converge to it over time, unstable (+) if nearby trajectories diverge from it, or a saddle (*) if nearby trajectories approach it in some direction(s) but diverge from it in some other direction(s). A fixed point is said to be a spiral (◦) if trajectories near the fixed point rotate either towards the fixed point (damped oscillation) or away from the fixed point (sustained oscillation or limit cycle in the present case). Strong oscillation mainly appears on the ascending branch of the red nullcline. Overall, we see that local connectivity defines the dynamics in each regime essentially by controlling the geometry of the nullclines.

### S5 The effects of nonlinearity in the local model

Before getting into the global model, we briefly demonstrate numerically how the present unified model (equation 1–2) extends the reduced Wong-Wang model [5, 6] to more complex scenarios. As expected, the dynamics of the present model match that of the reduced Wong-Wang model for low levels of excitation, i.e. weak local excitatory connectivity (Figure S3a-b). In a regime of stronger local excitatory connectivity, as explored in [7], the two models diverge (Figure S3c-d). In the present model (Figure S3c), all trajectories are well-confined within a physiologically plausible range—state variables *S_E_* and *S_I_* denote the fraction of open channels, which by definition are between 0 and 1. In contrast, certain trajectories of the reduced Wong-Wang model (Figure S3d) overshoot beyond the physiologically plausible range. The effect of added nonlinearity in the present model manifests through the curvature of the blue nullclines, which confines the flow of oscillatory activities and creates extended multistability (see e.g. Figure S2b). Thus, the present model is more suitable for studying key nonlinear dynamical features in the resting brain.

**Figure S3:**
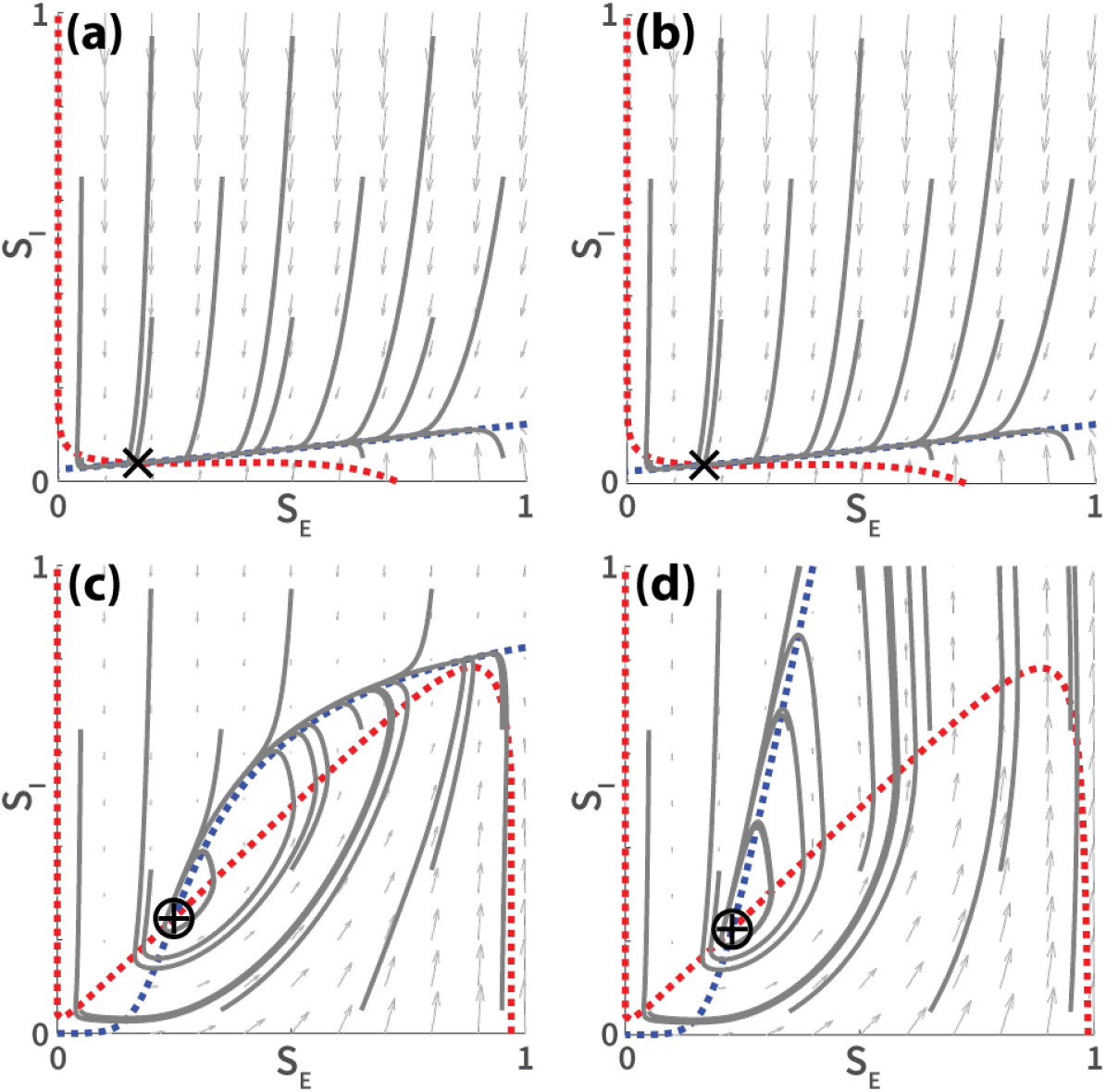
Comparisons between the present unified model (a, c) and the reduced Wong-Wang model (b, d) in two dynamic regimes. (a) and (b) show the phase portraits of the present model (equations 1–2) and the reduced Wong-Wang model (equations S4–S5) respectively in a regime of weak local excitatory connectivity. Parameter values are obtained from [6] and identical across the two models: *w_EE_* = 0.21, *w_EI_* = 0.15, *w_IE_* = 1, *w_II_* = 1, *I_E_* = 0.382 and *I_I_* = 0.267 (unspecified parameters follow Table 1). The resulted dynamics are virtually identical. (c) and (d) show a similar comparison between the two models in an oscillatory regime, where the local excitatory connectivity is stronger (*w_EE_* = 4, *w_EI_* = 1). While the dynamics of the present model (c) is well confined within a realistic range (*S_E_*, *S_I_* ∈ [0,1]), it is not the case for the reduced Wong-Wang model (d).

### S6 Local and global causes of temporal diversity

Now we turn to the structural constraints on temporal diversity. In particular, we show how spectral properties of the simulated neural activities and corresponding hemodynamic responses are affected by both the diversity of local structural properties and the structure of the large-scale connectome.

Given a uniform global network, temporal diversity across the whole brain can be induced by the diversity of local excitatory-to-excitatory connection (*w_EE_*), as shown in Figure S4a. Brain regions with relatively weak *w_EE_* (blue) have low characteristic frequencies around 10 Hz (alpha range), while brain regions with strong *w_EE_* (red) have higher characteristic frequencies around 30 Hz (beta/gamma range). In other words, the characteristic frequency of the oscillation increases monotonically with *w_EE_* (see also Figure S8a). This is expected from the behavior of isolated brain regions (Figure S1d). In addition to the expected diversity, signs of coordination between regions can be seen as the wide-spread alpha peaks (Figure S4a). In contrast, regions with a higher characteristic frequency (beta/gamma range) are not as influential to other regions. That is, low-frequency oscillations, rather than high-frequency ones, are responsible for global coordination.

The above observations concern high-frequency dynamics typically measured using, e.g. electroencephalography (EEG) and magnetoencephalography (MEG). For low-frequency dynamics typical for functional magnetic resonance imaging (fMRI), we examine the low-frequency content (0.01-0.1 Hz) of the normalized power spectra of BOLD activities, derived from the same simulated neural dynamics (see Section S7 in Supplementary Materials for details). The result is shown in Figure S4b: there is no significant dependency of low-frequency power on *w_EE_* (Spearman correlation *ρ* = −0.029, p = 0.81). In short, we find differential effects of local structural diversity on neural dynamics at the time scales typical for different neural imaging modalities.

**Figure S4:**
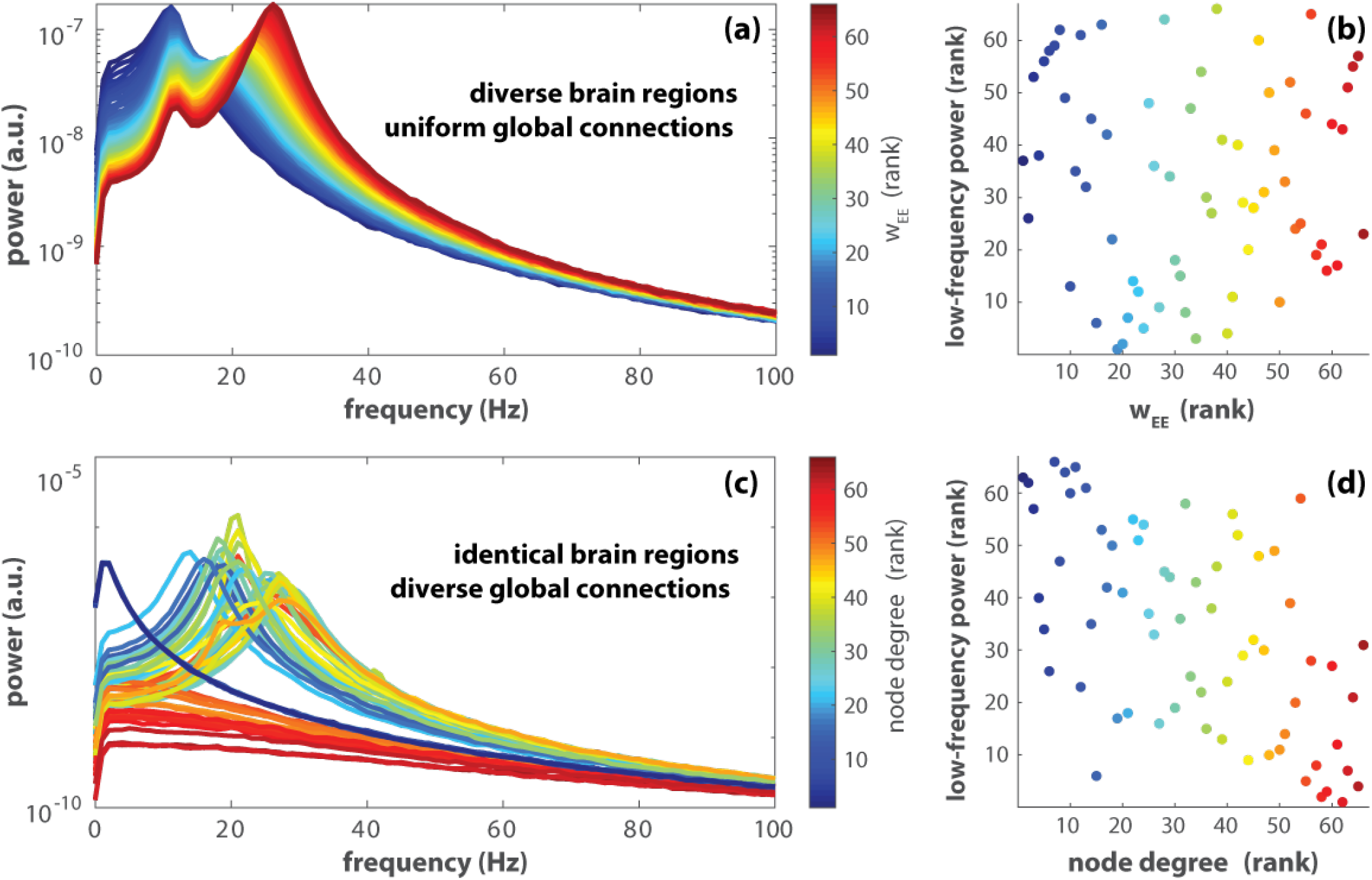
Temporal diversity induced by diversity in local (a, b) and global structural connectivity (c, d). Spectral analyses are based on two simulated trials of the global model (equation 4–6 with *N* = 66) each with identical initial conditions 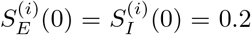, a 1200s duration, and a moderate level of noise *σ* = 0.01. For the first simulated trial (a, b), different brain areas are endowed with different local connectivity, 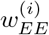, evenly spread in the interval [1, 2]; the large-scale structural connectivity is set to be uniform, i.e. *C_ij_* = 1/(*N* – 1), for *i* ≠ *j*. In addition, the global coupling *G* = 1.35. (a) shows the power spectra of the excitatory gating variables 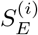 for *i* = 1,…, *N*. The spectrum for each brain region is color coded by the rank of *w_EE_*—blue to red indicate the smallest to the largest *w_EE_*. The peak frequency of these spectra clearly increases with *w_EE_*. (b) shows the rank of the low-frequency power of the corresponding BOLD signal, integrated over the frequency range [0.01, 0.1] Hz (see Section S7 for details), which depends little on the rank of *w_EE_*. (c) and (d) show results of similar analyses but for the second simulated trial, where the individual brain regions are identical (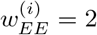 for all *i*) but the global structural connectivity is realistic, i.e. *C_ij_* here reflects the human connectome [11, 12] with the global coupling *G* = 2.5. Both low-frequency (d) and high-frequency (c) activities are highly affected by the degree of the brain region in the global network (rank color-coded).

On the other hand, temporal differentiation does not mandate the brain regions themselves to be structurally different. As shown in Figure S4c-d, locally identical brain regions can behave very differently due to the topology of the large-scale network (human connectome as in Section 2.2). The influence of large-scale structural connectivity on temporal diversity is manifested in both the high-frequency neural dynamics (Figure S4c; Figure S9a) and the low-frequency power of the BOLD signals (Figure S4d; Figure S9b). Specifically, the low-frequency power is inversely related to the degree of each node (brain region) in the large-scale network (Spearman correlation *ρ* = −0.584, *p* < 10^−6^).

In Section S12, we demonstrate that the above effects are robust over 200 simulated trials of the same parameter settings. Overall, both local (Figure S4a,b; Figure S8) and large-scale structural connectivity (Figure S4c,d; Figure S9) contribute to the diversification of local dynamics. The contribution of local structural differences is stonger in a higher-frequency range (Figure S4d; Figure S8a), while the contribution of global structural connectivity is stronger in a very-low frequency range (Figure S4d; Figure S9b). Modeling real neural dynamics requires considering both sides of the spectrum.

### S7 Computation of BOLD signal and low-frequency power

In the present study, we are interested in not only the high-frequency activity measurable by, for example, EEG recordings but also low-frequency fluctuations that are often a subject of investigation in fMRI studies. Therefore, we simulated the BOLD activities induced by the underlying neural dynamics and examine their low-frequency properties.

BOLD (Blood-oxygen-level-dependent) activities are computed using the Balloon-Windkessel model [13–16]. The hemodynamic response of the *i^th^* brain area takes the form

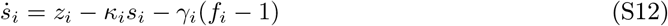

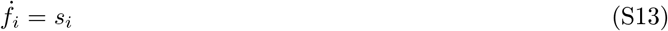

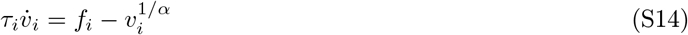

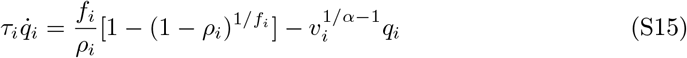

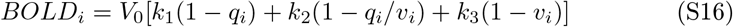

where the interpretation and value of the parameters are given in Table S3. The initial condition is

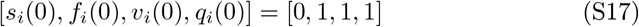

which is a hemodynamic equilibrium state without neural activity. *z_i_*(*t*) is the simulated neural activity, corresponding to the gating variable of the excitatory populations 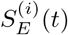.

**Table S3:** Parameters of the Balloon-Windkessel model of BOLD activities, obtained from [16].

| parameter | interpretation | value |
| --- | --- | --- |
| $z_i$ | neuronal activity | $S_E^{(i)}$ |
| $s_i$ | vasodilatory signal | variable |
| $f_i$ | blood inflow | variable |
| $v_i$ | blood volume | variable |
| $q_i$ | deoxyhemoglobin content | variable |
| $\kappa_i$ | rate of signal decay | $0.65(s^{-1})$ |
| $\gamma_i$ | rate of flow-dependent elimination | $0.41(s^{-1})$ |
| $\tau_i$ | hemodynamic transit time | $0.98 \text{ (s)}$ |
| $\alpha$ | Grubb's exponent | $0.32$ |
| $\rho$ | resting oxygen extraction fraction | $0.34$ |
| $V_0$ | resting blood volume fraction | $0.02$ |
| $k_1$ | BOLD weight parameter | $7\rho_i$ |
| $k_2$ | BOLD weight parameter | $2$ |
| $k_3$ | BOLD weight parameter | $2\rho_i - 0.2$ |

The power spectrum for each simulated BOLD time series is computed using Welch’s method [17], after being subsampled at 720ms intervals (matching the TR of resting state fMRI used in the Human Connectome Project [11]). The full power spectrum P (ω) was first normalized such that

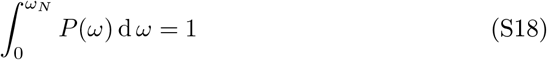

where *ω_N_* is the Nyquist frequency (approximately 0.7 Hz for the chosen sampling interval). The low-frequency power is defined as

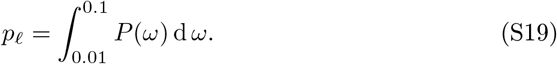

### S8 Discretization of regional states

Although the dynamic landscape of the global model can be quite complex (Figure 3g-i), each region in the globally connected network still falls into discrete states (Figure S5) very much like in the local model (disconnected stripes in Figure 3a-c)— it is the combination of regional states that produces a great variety of attractors at the global level. Discretized regional states (number on black disks in Figure S5) thus give rise to discretized attractors in the global model. Rank correlation (Spearman) between regional states across these discretized attractors are used to quantify *cross-attractor coordination* in the model brain (Figure 4b in the main text). Using discretized attractors, we quantify how much two regions move up and down together across attractors without considering the distance between the attractors. The distance between attractors are considered separately as the energy cost that constrains such transitions (Figure 5 in the main text).

**Figure S5:**
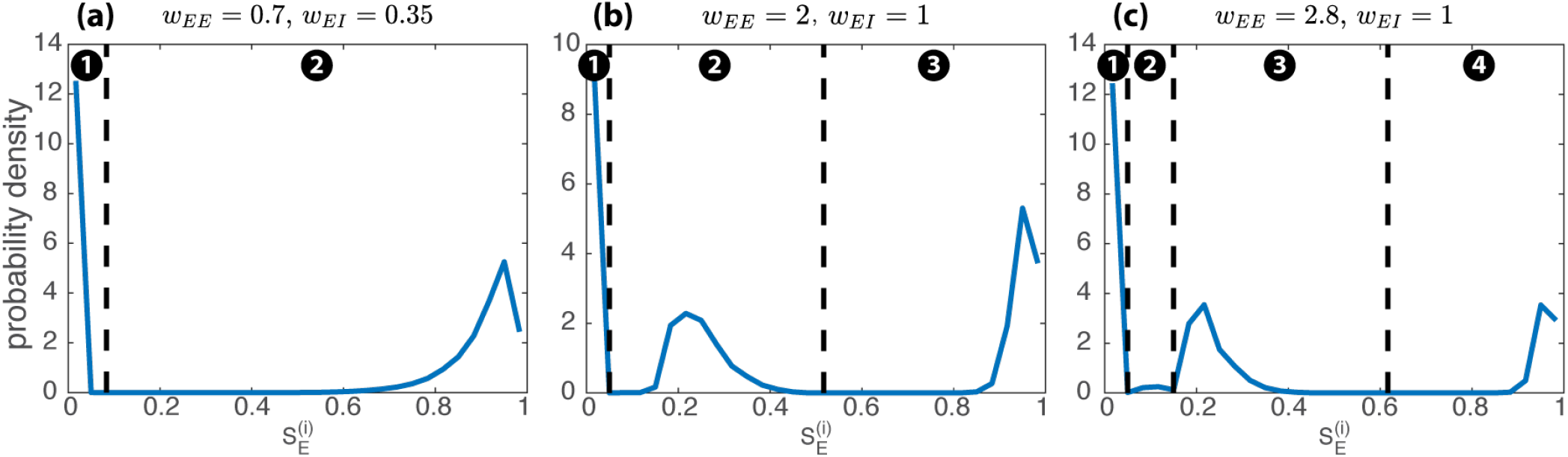
Brain regions fall into discrete states in the global model. Blue curves in (a-c) show the distribution of the state of individual brain regions (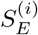 for any *i*) in the attractors in the bifurcation diagrams (Figure 3g-i) respectively. Black dashed lines indicate the location of local minima in the distributions. These minima are used to discretize the continuous variable 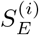 into integer-indexed (discrete) states (number in black disks).

### S9 Reliability of the average functional connectivity

The average functional connectivity is highly similar between Day 1 (Figure 4a in the main text) and Day 2 (Figure S6a). Linear regression analysis indicates that the Day-1 matrix is highly predictive of the Day-2 matrix (red x in Figure S6b; *β*_1_ = 0.98, t(2143)=203, p<0.001, *R*^2^ = 0.951; only elements below the diagonal are compared due to the symmetry of the matrix). Moreover, the functional connectivity matrix (Figure 4a in the main text) obtained using Spearman correlation is highly predictive of the corresponding Pearson correlation coefficients (Figure S6c; *β*_1_ = 1.03, t(2143)=897, p<0.001, *R*^2^ = 0.997), which itself is highly consistent across two days (blue o in S6b; *β*_1_ = 0.97, t(2143)=211, p<0.001, *R*^2^ = 0.954).

**Figure S6:**
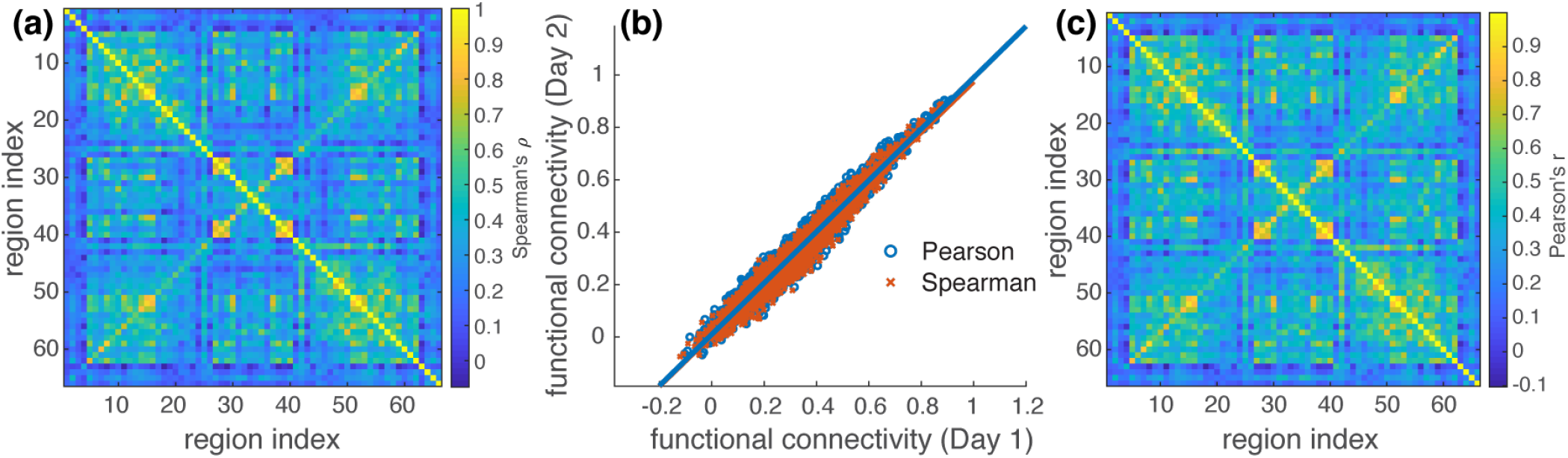
Average human functional connectivity highly reliable across two days and the types of correlation analysis. (a) shows the functional connectivity averaged across the same subjects as in Figure 4a (main text) but using the resting fMRI data (two runs) from Day 2. The matrices from the two days are highly correlated (b; red x). (c) shows the functional connectivity estimated using Pearson correlation averaged over all runs in Day 1, which is also consistent across two days (b; blue o), and is not markedly different from Figure 4a (main text). See text for statistical information.

### S10 Mean comparisons of human-model similarity

**Figure S7:**
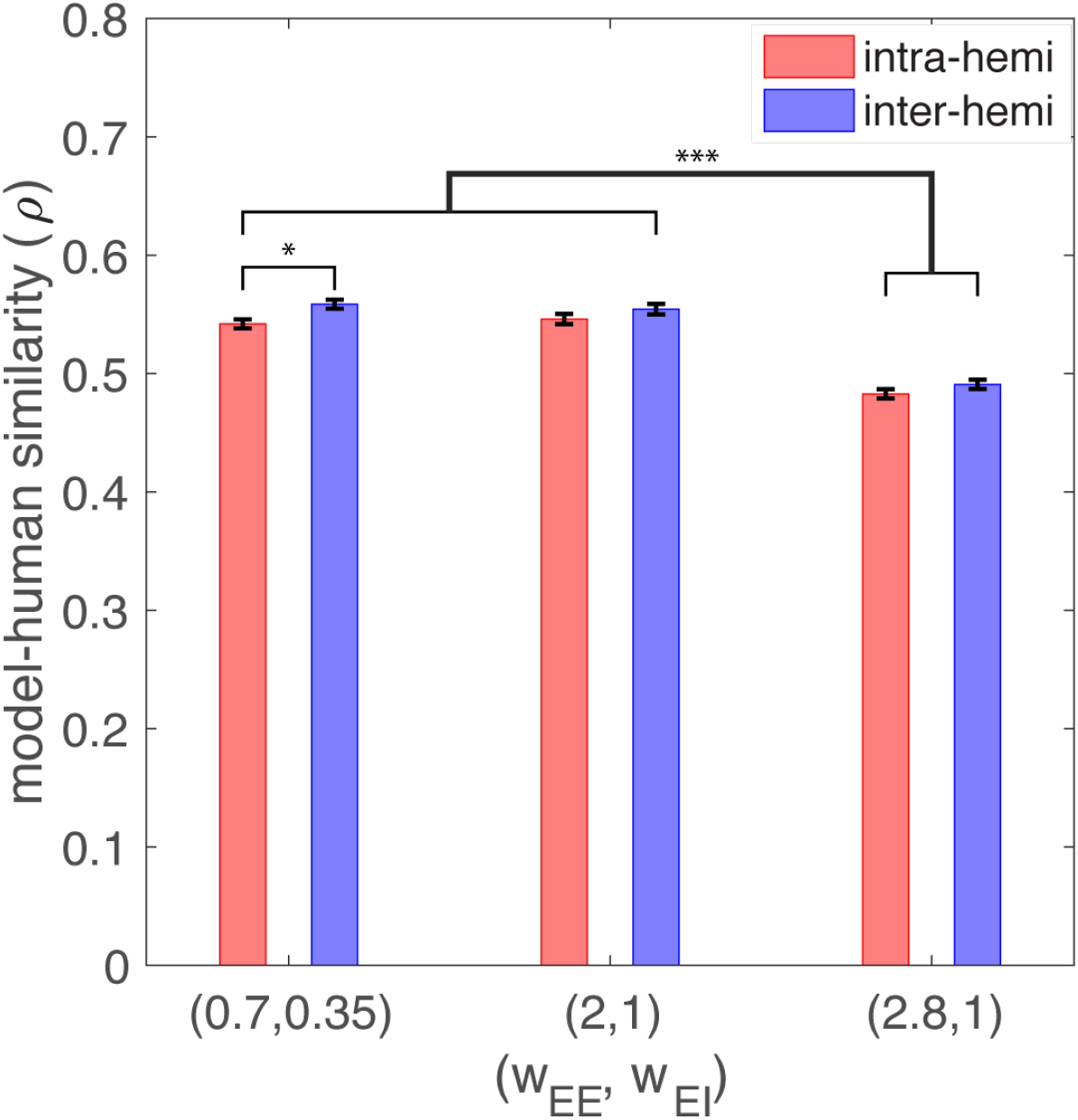
Average effects of local structural connectivity on model-human similarity. Cross-attractor coordination in the model well captures human functional connectivity for both intra-(red) and inter-hemispheric interaction (blue). The model-human correlation (Spearman’s *ρ*) is slightly worse when local excitatory connection is too strong (*w_EE_* = 2.8, *w_EI_* = 1). (* p<0.05, *** p<0.001, with Tukey HSD)

### S11 Results of permutation tests and bootstrapping

**Table S4:** Permutation tests for the loss of model-human similarity (Δ*ρ*/*ρ_all_*) by three different levels of local excitatory connectivity (*w_EE_*, *w_EI_*). The mean comparisons between different levels of *w_EE_*, *w_EI_* (bars in Figure 6) are based on 30000 random permutations. The significance levels are Bonferroni-corrected. A comparison reaches statistical significance if the mean is outside of the confidence interval.

| comparison | mean | $\alpha$ | confidence interval |
| --- | --- | --- | --- |
| $loss(0.7, 0.35) - loss(2, 1)$ | 0.1777 | 0.001 | (-0.1036, 0.0999) |
| $loss(0.7, 0.35) - loss(2.8, 1)$ | 0.3410 | 0.001 | (-0.1011, 0.1030) |
| $loss(2, 1) - loss(2.8, 1)$ | 0.1633 | 0.001 | (-0.0993, 0.0980) |

**Table S5:**
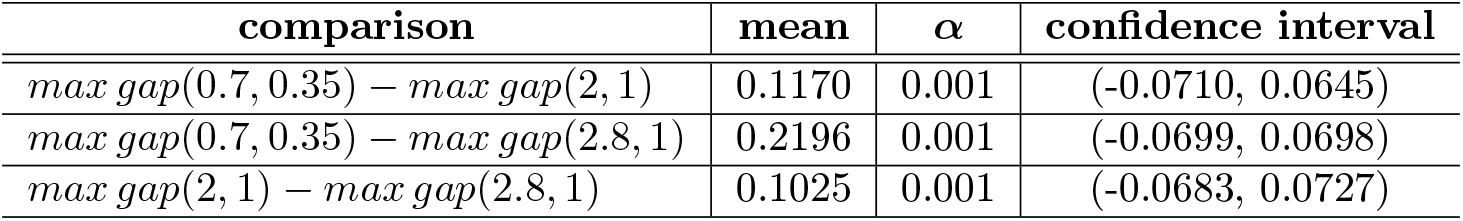
Permutation tests for the maximum energy gaps by three different levels of local excitatory connectivity (*w_EE_*, *w_EI_*). The mean comparisons between different levels of *w_EE_*, *w_EI_* (bars in Figure 5b) are based on 30000 random permutations. The significance levels are Bonferroni-corrected. A comparison reaches statistical significance if the mean is outside of the confidence interval.

**Table S6:** Permutation tests for the mean energy gaps by three different levels of local excitatory connectivity (*w_EE_*, *w_EI_*). The mean comparisons between different levels of *w_EE_*, *w_EI_* (bars in Figure 5c) are based on 30000 random permutations. The significance levels are Bonferroni-corrected. A comparison reaches statistical significance if the mean is outside of the confidence interval.

| comparison | mean | $\alpha$ | confidence interval |
| --- | --- | --- | --- |
| $\text{mean gap}(0.7, 0.35) - \text{mean gap}(2, 1)$ | 0.0405 | 0.001 | (-0.0084, 0.0085) |
| $\text{mean gap}(0.7, 0.35) - \text{mean gap}(2.8, 1)$ | 0.0496 | 0.001 | (-0.0089, 0.0084) |
| $\text{mean gap}(2, 1) - \text{mean gap}(2.8, 1)$ | 0.0090 | 0.001 | (-0.0083, 0.0087) |

### S12 Dependency of spectral properties on local and global structural connectivity

In Figure S4, we illustrate with two simulated trials how high-frequency and low-frequency dynamics depend on local excitatory-to-excitatory connectivity *w_EE_* and the topology of the global network. To show that these effects are not incidental, we simulated 200 trials for each of the conditions: (1) the global network is uniform but local connectivity *w_EE_* is diverse (as in Figure S4a,b), and (2) local connectivity *w_EE_* is identical but the global network follows the human connectome (as in Figure S4c,d). We characterize the high-frequency content of a spectrum as its peak frequency, i.e. the frequency at which the spectral power is the highest (e.g. peaks in Figure S4a,c); the low-frequency content as the integral of the power between 0.01 and 0.1 Hz (Section S7). The dependency of these features on local (*w_EE_*) and global structural properties (node degree) is quantified using Spearman correlation. The distributions of the correlation coefficients (*ρ*) and corresponding p-values are shown in Figure S8 and Figure S9 for condition 1 and 2 respectively. Figure S8 shows that local structural connectivity *w_EE_* strongly affects the peak frequency of the brain region (a,c) but not so much the low-frequency power (b,d). The stronger the local connectivity, the higher the peak frequency. Figure S9 shows that the node degree of the global network has a strong and negative effect on the low-frequency power, and a weak and positive effect on the peak frequency. Figure S4 illustrates such dependencies using typical trials (median correlation coefficients) from the distributions (Figure S8–S9).

**Figure S8:**
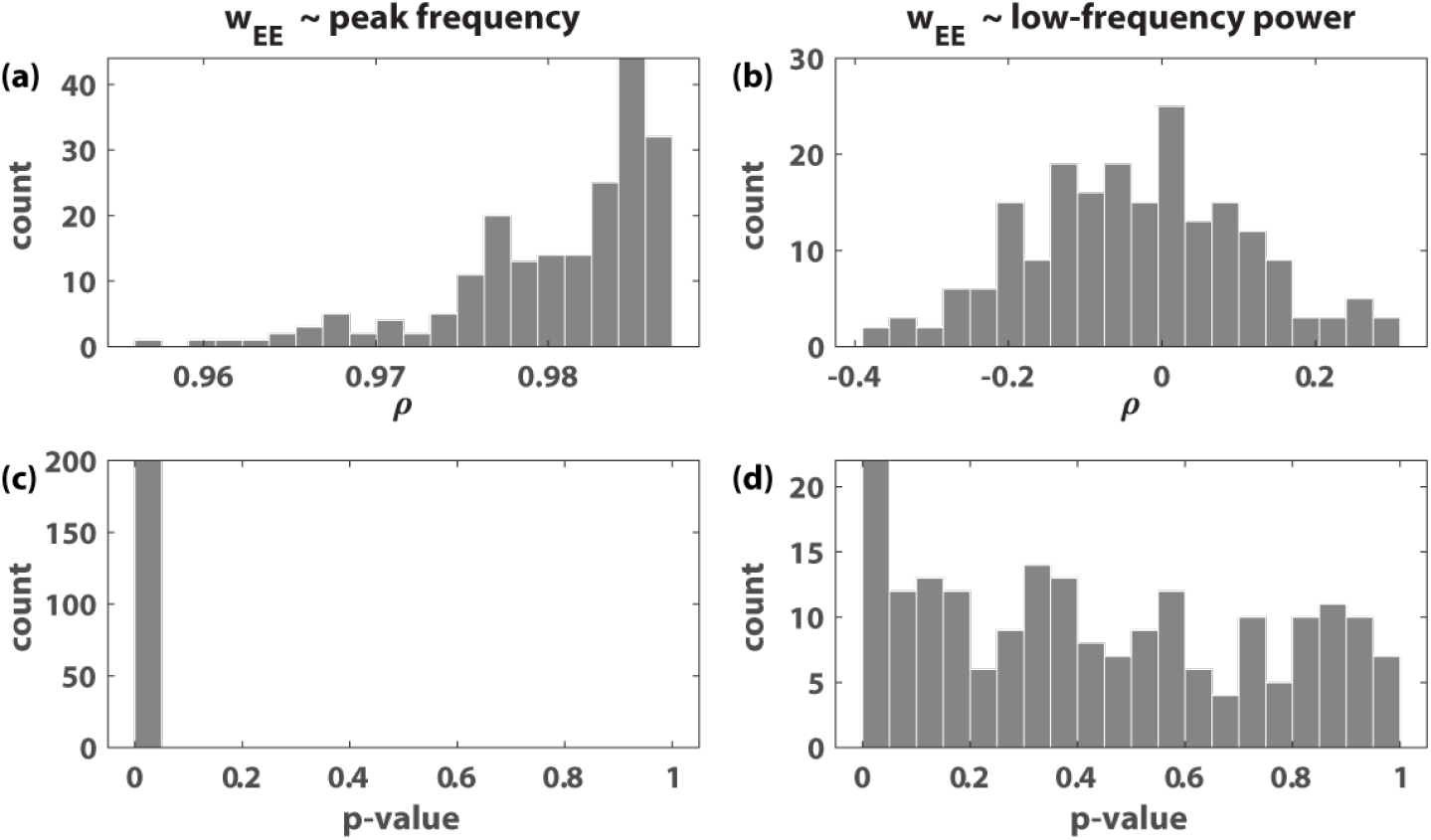
Dependency of peak frequency and low-frequency power on local excitatory-to-excitatory connectivity. 200 trials are simulated following the same parameter setting as Figure S4a,b, where the global network is uniform but the local connectivity *w_EE_*’s spread between 1 to 2 for different brain regions. The noise terms in equation 4–5 make these trials different realizations of the same noisy process. The peak frequency of the spectra, e.g. from 10 to 30 Hz in Figure S4a, strongly depends on local connectivity *w_EE_* (a: *ρ*’s all close to 1; c: p-values all less than 0.05). In contrast, low-frequency power does not significantly depend on *w_EE_* (b: *ρ*’s distribute around zero; d: p-values spread between 0 and 1). Figure S4b shows this lack of dependency in an example trial that corresponds to the median of the distribution (b).

**Figure S9:**
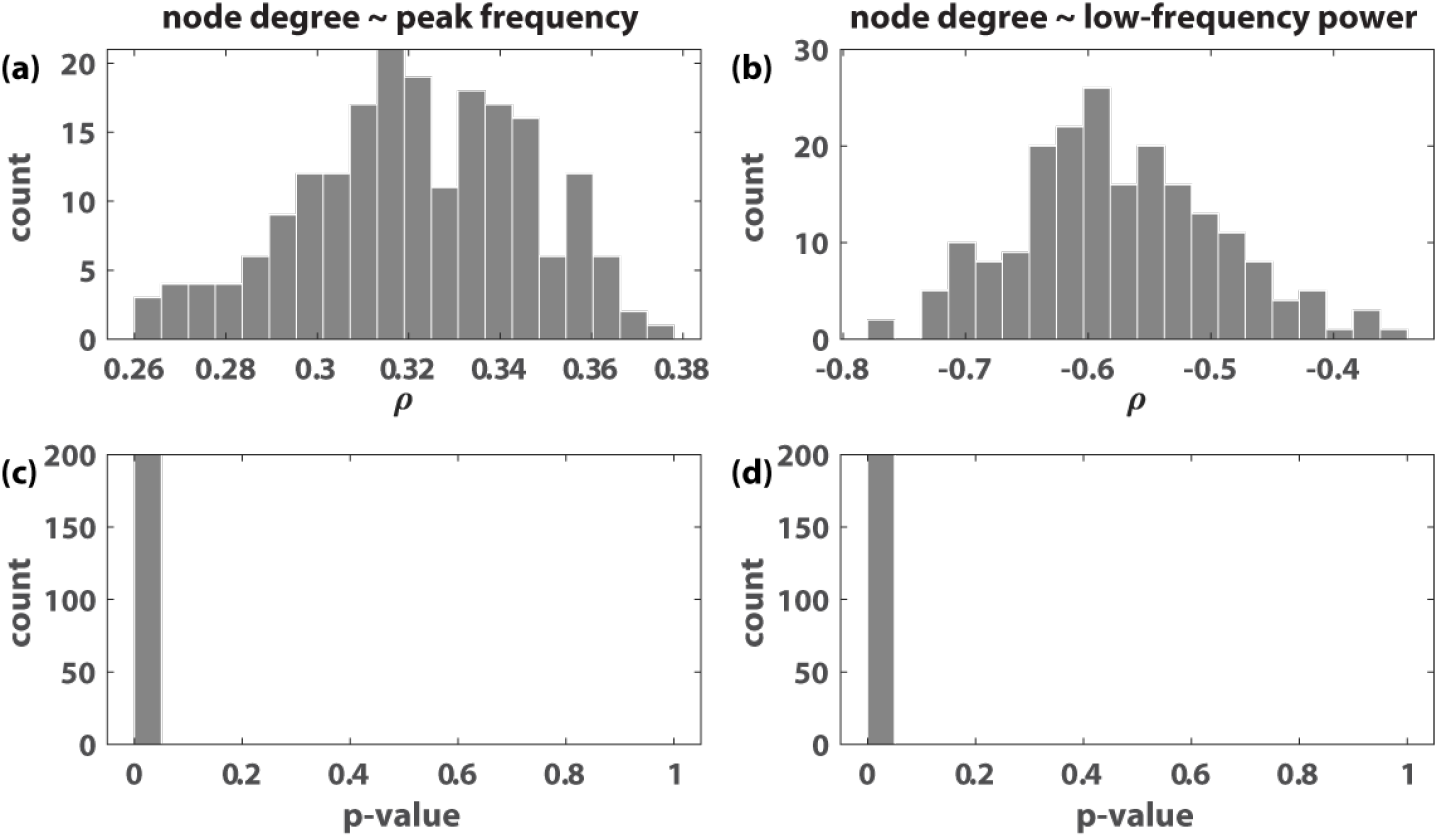
Dependency of peak frequency and low-frequency power on node degree in the global network. 200 trials are simulated following the same parameter setting as Figure S4c,d, where the local connectivity *w_EE_*’s are the same across brain regions but the global-network reflects the human connectome (see main text). The peak frequency of the spectra, e.g. between 0 and 30 Hz in Figure S4c, moderately increases with the node degree of each region (a: positive *ρ*’s around 0.32; c: p-values all less than 0.05). Low-frequency power decreases more significantly with node degree (b: *ρ*’s distribute around −0.6; d: p-values all less than 0.05). Figure S4d illustrates this dependency with an example trial that corresponds to the median of the distribution (b).

### S13 Analysis of the local model

We can see from the numerical analysis that the nullclines (dashed lines in Figure S2) crucially constraint the dynamics of the local model (equation 1–3). Here we take a closer look at their shapes. Red nullcline indicates where there is only vertical flow,

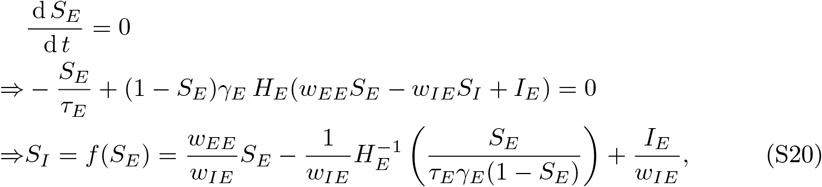

and blue nullcline indicates where there is only horizontal flow,

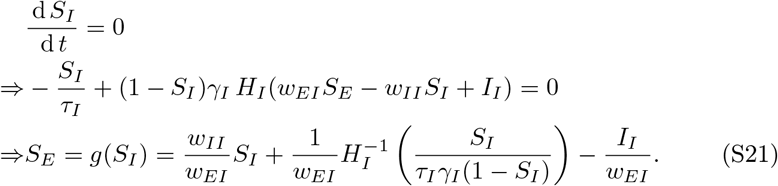

What is common between the two nullcines, *S_I_* = *f*(*S_E_*) and *S_E_* = *g*(*S_I_*), is that their shape crucially depends on a linear term *S_p_* and the inverse of the transfer function 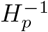 for *p* ∈ {*E*, *I*}. Both terms are monotonically increasing with *S_p_* (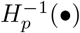 and *S_p_*/(1 – *S_p_*) are both monotonically increasing function; so is their composition). 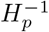 is only defined on a domain between 0 and *r_max_*, for which the nullclines are confined within the interval

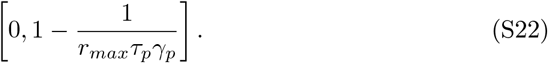

Within this interval *S_I_* = *f*(*S_E_*) (equation S20; red nullcline), overall, goes down from +∞ to −∞, while *S_E_* = *g*(*S_I_*) (equation S21; blue nullcline) goes up from −∞ to +∞. This results from the dominant effect of 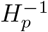 for a very large or very small input.

In between these extremes, the effect of the linear term is more pronounced. This is especially the case for *S_I_* = *f*(*S_E_*) (red nullcline): the linear term monotonically increases with *S_E_*, counteracting the descending trend of 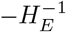. Given a sufficiently strong excitatory-to-excitatory connection *w_EE_* (self-excitation), the linear term “twists” the nullcline counterclockwise, creating an ascending branch in the middle. If we balance the level of self-excitation with inhibitory-feedback—let *w_EE_* = *w_IE_*—equation S20 becomes

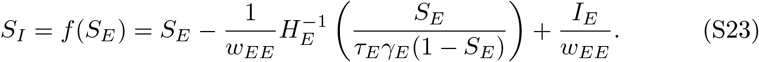

In this simplified case, increasing self-excitation *w_EE_* reduces the influence of 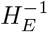 such that the slope of middle branch approaches 1.

For *S_E_* = *g*(*S_I_*) (equation S21), the linear term and the 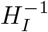 term increase together, so that *S_E_* = *g*(*S_I_*) (blue nullcline) is always monotonically increasing. Given a fixed *w_II_*, *S_E_* = *g*(*S_I_*) increases with *S_I_* at an overall slower rate for larger *w_EI_*, or more conveniently seen as *S_I_* = *g*^−1^(*S_E_*) increasing faster with *S_E_* for larger w_EI_. Intuitively, increasing *w_EI_* twists *S_E_* = *g*(*S_I_*) counterclockwise, seen as the middle segment of the blue nullcline becoming more vertical.

We have discussed above how local connectivity *w_EE_* and *w_EI_* influence the gross geometry of the nullclines—twisting the middle segment of the curve counterclockwise. But how are these geometric changes going to affect the dynamics? We show below that they critically control the multistability and oscillation in the local model.

#### Multistability

For the local model to be multistable, *S_I_* = *f*(*S_E_*) (red nullcline) must have an ascending branch, i.e. f (*S_E_*) cannot be monotonically decreasing.

##### Proof.

Suppose that *f*(*x*) and *g*^−1^(*x*) are monotonic functions for *x* ∈ [0, 1]. Specially, *g*^−1^(*x*) is monotonically increasing; *f*(*x*) is monotonically decreasing. Assume that *f*(*x*) and *g*^−1^(*x*) intersect at two points *x*_1_ ⩽ *x*_2_, i.e. *f*(*x*_1_) = *g*^−1^(*x*_1_) and *f*(*x*_2_) = *g*^−1^(*x*_2_). Since *g*^−1^(*x*) is monotonically increasing, we have *g*^−1^(*x*_1_) ⩽ *g*^−1^(*x*_2_), which implies *f*(*x*_1_) ⩽ *f*(*x*_2_). Meanwhile, since *f*(*x*) is monotonically decreasing, *f*(*x*_1_) ⩾ *f*(*x*_2_). Thus, we have *f*(*x*_1_) = *f*(*x*_2_), and by monotonicity, *x*_1_ = *x*_2_. In other words, if the two functions intersect, there must be a unique intersection.

Since *g*^−1^(*x*) is always monotonically increasing and the existence of multistability requires the existence of multiple intersections between *g*^−1^(*x*) and *f*(*x*), a monotonically decreasing *f*(*x*) implies that the system cannot be multistable. In other words, if the system is multistable, then *f*(*x*) cannot be monotonically decreasing.

This result highlights the importance of self-excitation *w_EE_* in equation S20 and equation S23-multistability can only occur when *w_EE_* is sufficiently large. Correspondingly in the numerical result (Figure S1), the region of multistability appears only for larger *w_EE_*’s.

Note that the above argument is not restricted to the present model, but applicable to models that share the geometry form of the Wilson-Cowan model in general. Nevertheless, one would hope to know how large a *w_EE_* is large enough for multistability to be possible, and this depends on the specific formulation of the transfer function (equation 3) and the underlying assumptions about neuronal level properties (equation S8). Ideally, to know the minimal *w_EE_*, one need to find the minimal slope of 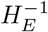 (*u*(*S_E_*)) with respective to *S_E_*, where *u*(*S_E_*):= *S_E_*/(*τ_E_γ_E_*(1 – *S_E_*)). The exact solution is, however, rather perplexing to calculate. Here we provide a rough, but simple, estimation instead. The slope of interest is

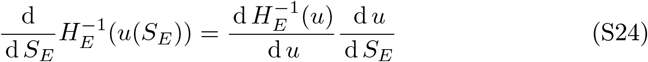

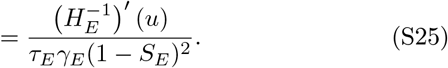

Instead of finding the minimum of equation S25, we aim to find a representative point 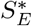 such that equation S25 is relatively small.

One option is to use the minimum of the numerator. The minimum of the numerator 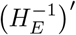 (*u*) is simply the reciprocal of the maximum of 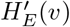, where 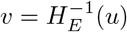. By design, *H_E_* reaches its maximum slope *a_E_* at the inflection point 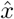, where *H_E_*(*v*) = *r_max_*/2. That is, we need

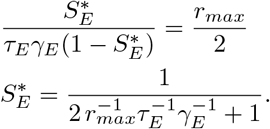

But note here that, in the case where *r_max_* is a large number, the representative point 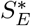 is very close to one, which further results in a small denominator in equation S25 and a large slope for 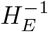. Thus, the inflection point of *H_E_*(*v*) is not a very good choice.

To avoid the small denominator problem for equation S25, we need to choose a 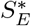 as small as possible while *H_E_*(*v*) remains close to the line *a_E_v* – *b_E_*. For this purpose, we take *v*\* to be the intersection between the line *a_E_ v* – *b_E_* and the horizontal axis,

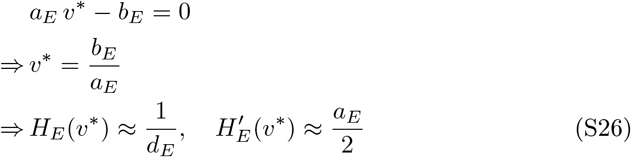

(approximate values can be obtained from the Taylor expansion of 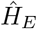 near *v*\*). Given equation S26, we need

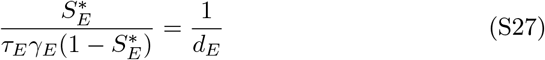

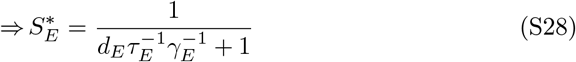

and

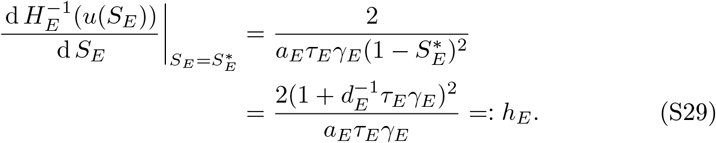

Now for the nullcline *S*_1_ = *f*(*S_E_*) to have a positive slope at 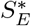, one simply needs

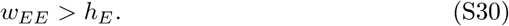

Here *h_E_* is approximately 0.2 based on the present parameter choices, inherited from Wong and Wang’s initial derivation [5]. This result is confirmed numerically by the bifurcation diagrams (Figure 3a-c vs. Figure S10a) of the local model—multistability exists for some level of input *I_E_* when *w_EE_* > 0.2.

#### Oscillation

Now we look for the conditions for oscillation to emerge. Here we are mainly concerned with the oscillation occurring on the ascending segment of *S_I_* = *f*(*S_E_*) (red nullcline). Following a similar argument as Wilson and Cowan [2], one notice that for the flow around a fixed point—an intersection between the nullclines—to have consistent rotation, the nullcline *g*^−1^(*S_E_*) (blue) must have a greater slope than *f*(*S_E_*) (red nullcline). Qualitatively, one would expect oscillation to be induced by increasing *w_EI_*, which twists *g*(*S_I_*) (blue nullcline) counterclockwise. This expectation is confirmed by the numerical results in Figure S1a-d: oscillation emerges for sufficiently large *w_EI_* for fixed points on the ascending branch of *S_I_* = *f*(*S_E_*) (Figure S2a-d).

Quantitatively, we consider the derivative of the two nullclines at a respective representative point. First, we extend the results in equation S28–S29 to the second nullcline *S_E_* = *g*(*S_I_*) (blue):

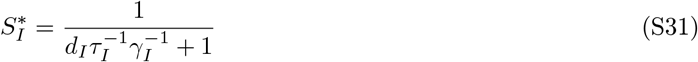

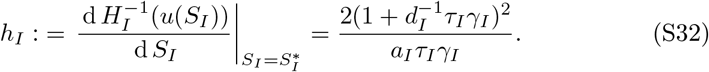

For parameters used in the present study, *h_I_* ≈ 0.4. We have the slope of the two nullclines at their respective representative points,

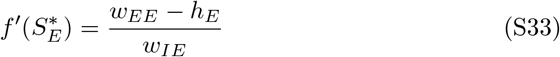

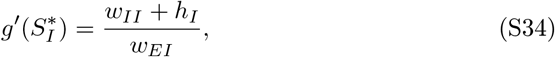

and we need

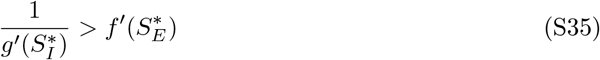

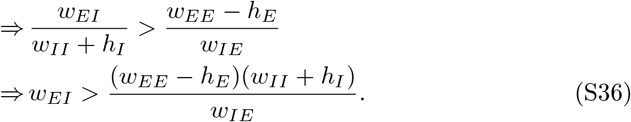

With balanced inhibitory feedback *w_IE_* = *w_EE_*, we have

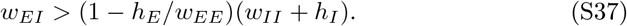

For very large *w_EE_*, one simply need

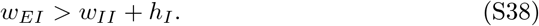

Given the present parameter choices, we need *w_EI_* > 0.45 to induce oscillation for some level of input *I_E_* and *I_I_*. This is in line with the numerical results in Figure S1. For *h_E_* > 0, as assumed here, lowering w_EE_ also lowers the threshold for oscillation.

#### Linear stability analysis

In addition to the presence of oscillation, one would also want to know if such oscillation is sustainable or damped. Here we extend the above analysis by linearizing the system near a specific fixed point. A fixed point is where the two nullclines (equations S20–S21) intersect. Conveniently, we let them intersect at their respective representative points 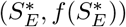 and 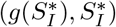 (see equation S28 and equation S31),

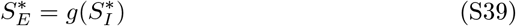

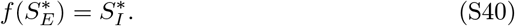

The two equations can be satisfied by the appropriate choice of *I_E_* and *I_I_*. The fixed point of our choice 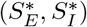 inherits a couple of properties from the above analysis, which we shall soon see. First, we define

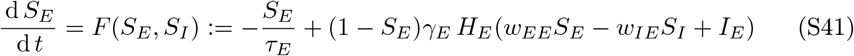

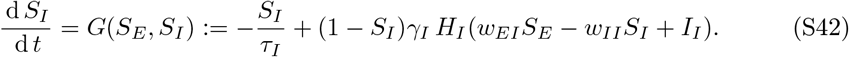

At the fixed points, we have from equation S41

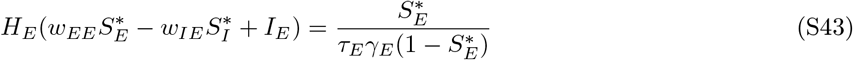

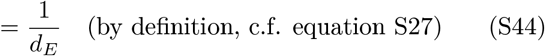

which implies that

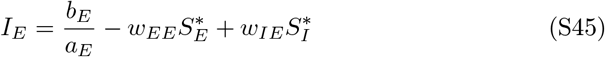

and

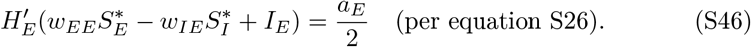

Similarly from equation S42, we have

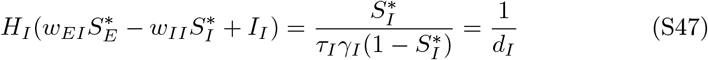

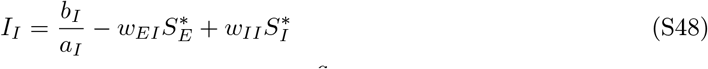

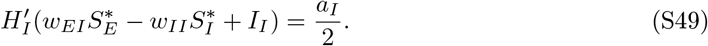

Now we are take the partial derivatives of *F* and *G* at 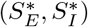,

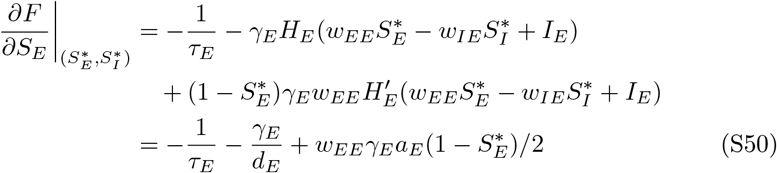

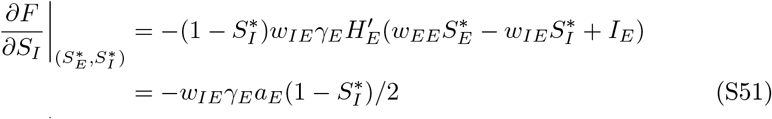

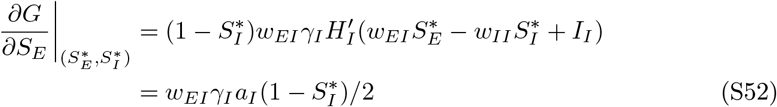

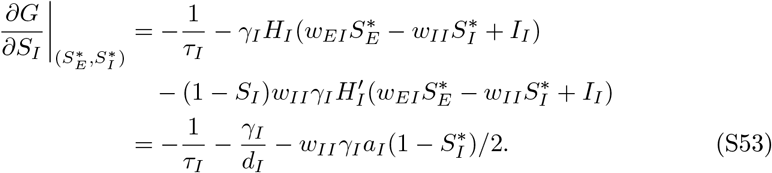

For simplicity, let parameters

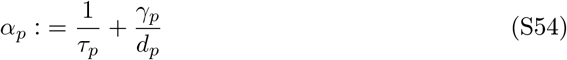

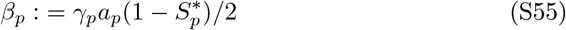

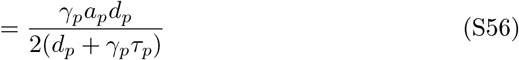

for *p* ∈ {*E*, *I*}. Note that by definition, both *α_p_* and *β_p_* are positive. Given parameters used in the present study, we have *α_E_* ≈ 14, *α_I_* ≈ 111, *β_E_* ≈ 71, and *β_I_* ≈ 276.

We write the Jacobian matrix as

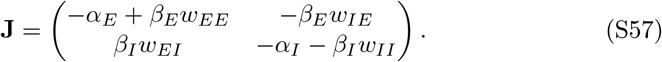

The eigenvalues of the Jacobian are

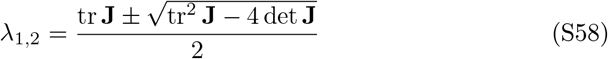

where

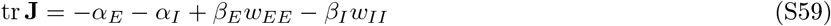

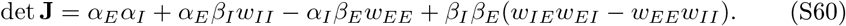

Assuming that the system is already oscillatory near the fixed point, i.e. tr^2^ **J** < 4 det **J**, to have sustained oscillation (limit cycle), we need

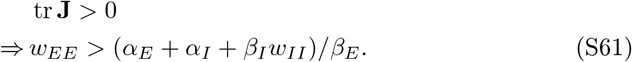

Given the parameters used in the present study, the emergence of limit cycles requires *w_EE_* > 2. Correspondingly in the numerical results (Figure S1), equation S61 provides an estimate of the lower bound of the Hopf bifurcation (gray dashed line). Note that stronger inhibitory-to-inhibitory connection *w_II_* increases the minimal *w_EE_* required to induce sustained oscillation. Overall, these analyses show that sustained oscillation requires both strong self-excitation and a sufficiently active inhibitory population.

In summary, we have shown analytically how structural connectivity *w_EE_* and *w_EI_* critically shape the dynamics—in this very low-dimensional parameter space, the system can easily switch between qualitatively different behavior. In particular, excitatory-to-inhibitory connectivity *w_EI_* controls the emergence of oscillation; excitatory-to-excitatory connectivity *w_EE_* controls both the emergence of multistability and sustained oscillation. The qualitative description of the system only depends on the gross geometric form of the Wilson-Cowan model, but the exact boundaries between regimes depend on the specific transfer function and the associated biophysical constraints.

### S14 Analysis of the global model

Now we take a look at the deterministic version of the global model,

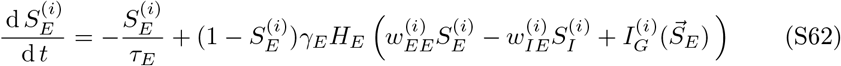

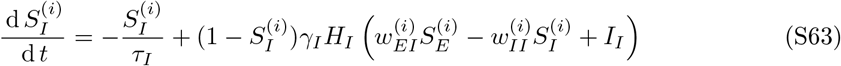

where

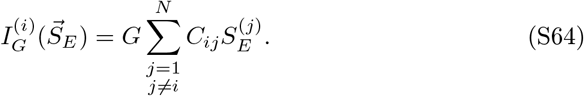

In this case, the nullclines are hyper-surfaces,

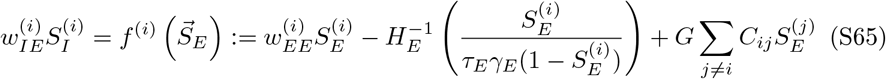

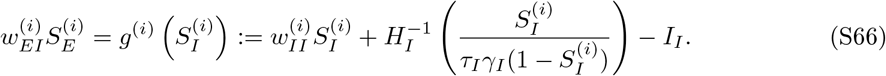

From equation S65 one can see that the local effect of global coupling is simply tilting the nullcline 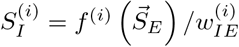 upwards with respect to 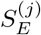.

The tilting of the nullcline 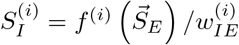 impact its number of intersections with 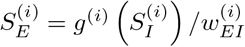 in each level set of 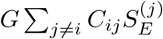. The number of intersections consequently constrains the number of stable states. A precise characterization of intersections is beyond the scope of the present work. Nevertheless, we hope to provide a few insights about the global geometry below.

#### Multistability

Following a similar argument as for the local model, we first show that, without global interaction (i.e. *G* = 0), the system cannot be multistable, if *w_EE_* is sufficiently small such that 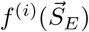 monotonically decreases with 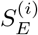 for all *i*. As shown above, the monotonicity condition implies that each local node by itself is not multistable.

##### Proof.

Assume there are at least two distinct fixed points of the system: 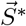 and 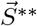, where 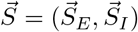 and 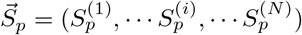 for *p* ∈ {*E*, *I*}. Since they are distinct points, there exists an 0 < *i* ⩽ *N* such that 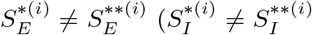 implies 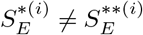 due to the monotonicity of *g*). Without loss of generality, we let 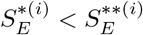.

Since we know that 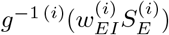 is always a monotonically increasing function, we have

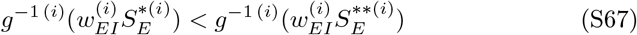

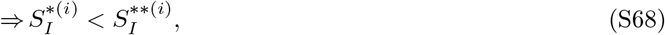

which also implies that

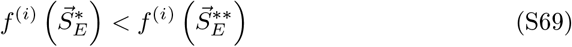

by definition of the nullcline 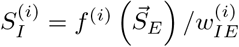, for any choice of *G* and *C_ij_*.

Now if 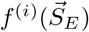 is monotonically decreasing with respect to 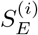, we know that at least for *G* = 0,

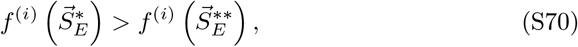

which leads to a contradiction. Thus, if the system has multiple fixed points, 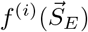 cannot be monotonically decreasing with respect to 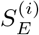 for all *i* when *G* = 0.

However, given a sufficiently large global coupling, especially for *G* > 1, multistability becomes possible.

##### Proof.

Following the above proof, the assumption 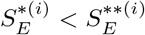 leads us to equation S69, or Δ_*G*_ < 0, where

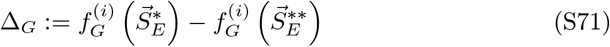

for any global coupling *G* ⩾ 0.

On the other hand, for the special case of *G* = 0, we can plug equation S65 into the definition S71 and have

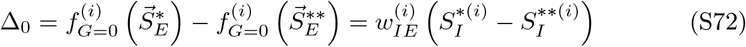

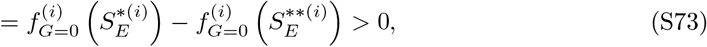

by our assumption that 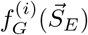 is a monotonically decreasing function with respect to 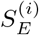. Since by definition, the coordinates of each fixed point is bounded between 0 and 1, we have

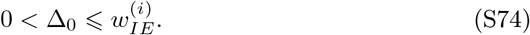

In the case of *G* = 0, this leads to a contradiction Δ*_G_* > 0, as we have already shown above.

Now we consider what happens when *G* > 0. Again, by plugging equation S65 into the definition S71, we have

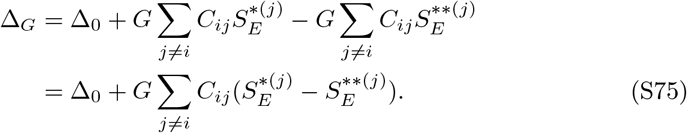

We need a bound on the second term in equation S75. Since *G* > 0 and *C_ij_* ⩾ 0,

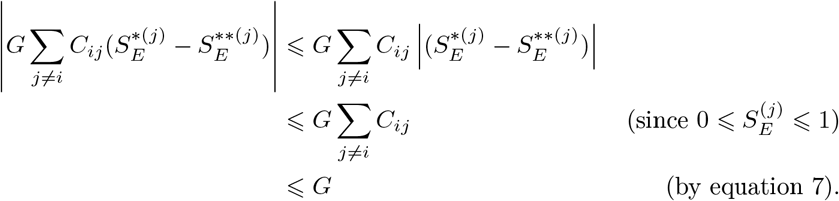

This gives us

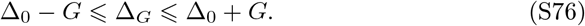

Thus, contradiction with equation S69 is inevitable if *G* < Δ_0_. On the other hand, by equation S74, we know that for *G* > Δ_0_, there exists some 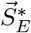 and 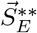 for some global network *C_ij_* such that Δ*_G_* < 0 consistent with equation S69. Thus it is possible for the global model to be multistable if *G* > Δ_0_, especially if 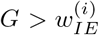, or 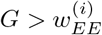 given matched inhibitory feedback *w_EE_* = *w_IE_*. This does not mean, however, that the system has to be multistable, due to the dependency on *C_ij_*.

To summarize, the above analyses suggest that a collection of brain regions that have no independent memory capacity (i.e. multistability) can *acquire* memory capacity when connected to each other in a global network, given sufficient global coupling. We further support this claim with numerical analysis (Figure S10). We refer to this kind of memory as *synergistic* memory—it is an emergent property that the parts themselves do not possess.

**Figure S10:**
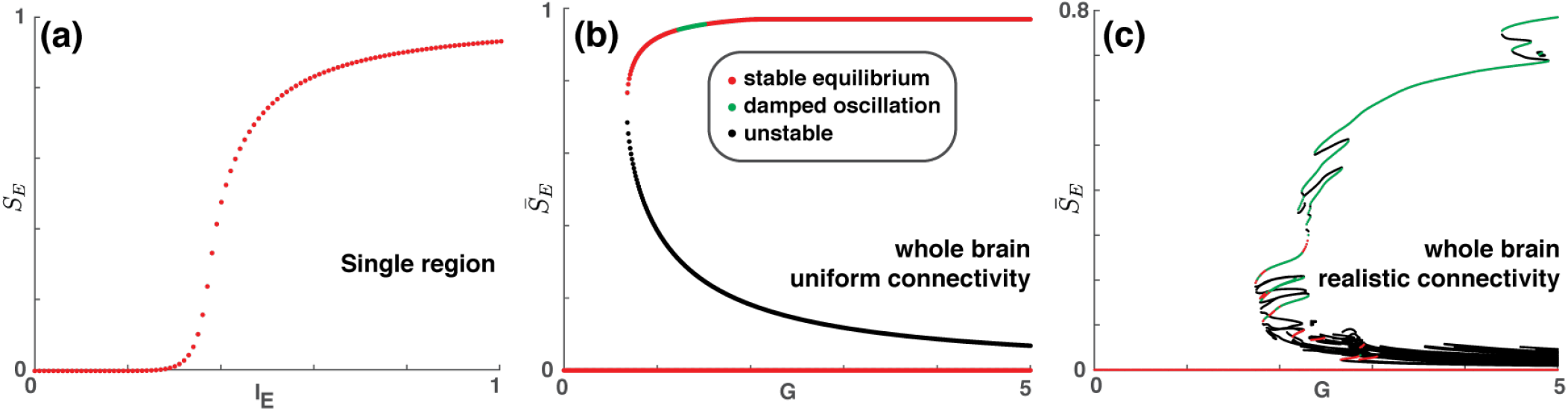
Synergistic memory between monostable nodes. Three bifurcation diagrams are shown for local parameters *w_EE_* = 0.1 and *w_EI_* = 0.35. They correspond to Figure 3a, d, g but with a lower *w_EE_* such that each local node by itself is monostable for any level of input (a). While each local node is completely monostable (no memory capacity), once there is sufficient global coupling *G* between them, the whole brain acquires memory capacity (b, c) that cannot be attributed to the parts alone—synergistic memory. Nevertheless, the size of the global memory capacity is still fundamentally constrained by the complexity of the local node (42 attractor branches in (c), very small compare to Figure 3g, h, i). See text for further discussion.

What we have not addressed in the above analyses is to what extent the global system is multistable—what is the number of stable states, or the size of the memory capacity—and what are the contributions from local self-excitation and global network connectivity. An analytical approach to this problem is difficult; thus, it is mainly addressed numerically (c.f. Figure S10 and Figure 3). Nevertheless, we provide an intuitive argument below as to how local and global connectivity affects the relevant geometrical properties of the dynamical system.

#### Local origin of geometrical complexity

At an intuitive level, the number of intersections between these hypersurfaces (equation S65–S66) is likely to increase with the number of folds of each surface. In the present case, the folding of hypersurfaces entails the temporary reversal of the sign of its partial derivative along a certain direction. Observe equation S65 and see that global coupling cannot create any folding of the surfaces. Thus, the geometrical complexity of the nullclines purely depends on the local properties of each node, in particular, the folding effect of self-excitation 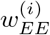.

#### The effect of global coupling

Without global coupling (*G* = 0), the number of fixed points of the global model is simply

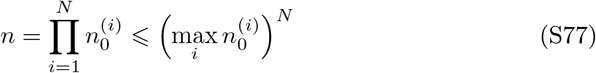

where 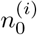 is the number of fixed points for each corresponding local model when *I_E_* = 0. Introducing global coupling (*G* ≠ 0) tilts each surface (equation S65) in a way dependent on the structure connectivity *C_ij_*. This may remove or introduce new intersections between the surfaces without changing the geometrical complexity of these surfaces. Thus, global coupling allows system-level multistability to be created synergistically, given appropriate structural connectivity *C_ij_*.

In summary, local and global coupling produce different geometrical effects on the system and jointly affect the number of possible stable states.

## Notes

### Competing Interest Statement

The authors have declared no competing interest.

### Summary of Updates

Adding additional validation results.

